# Neonatal Microglia and Their Secretome as Mediators of Brain Repair

**DOI:** 10.1101/2025.09.21.676806

**Authors:** Gabriela Lyszczarz, Eydís Sigurðardóttir Schiöth, Anouk Benmamar-Badel, Dominika Rusin, David Ramonet, Marco Anzalone, Thalissa Caers, Lejla Vahl Becirovic, Kirstine Nolling Jensen, Sanne G.S. Verberk, Fleur Mingneau, Martyna Makowska, Kate Lykke Lambertsen, Amy F. Lloyd, Jerome J. A. Hendriks, Agnieszka Wlodarczyk

## Abstract

Microglia are essential regulators of myelin integrity and repair, yet their regenerative capacity declines with ageing and in neurodegenerative diseases such as multiple sclerosis (MS). Neonatal microglia retain a uniquely reparative program that may offer insight into restoring lost functions in the adult CNS. Here we show that transplantation of neonatal microglia ameliorates disability, reduces leukocyte infiltration, and promotes remyelination in both inflammatory (EAE) and non-inflammatory (cuprizone) models, and reverses cognitive decline in aged mice. These benefits persisted even when transplanted cells remained confined to the meninges and were reproduced by the neonatal microglia secretome, indicating a paracrine mechanism. Multi-omic profiling revealed that the neonatal secretome is enriched in trophic factors and membrane-building lipids compared to adult microglia, while transcriptomic analyses of treated aged brains showed reactivation of developmental repair pathways and suppression of inflammatory signatures. Together, these results demonstrate that neonatal microglia re-engage rejuvenation-like programs in the adult CNS and highlight the importance of multifactorial strategies, integrating trophic, metabolic, and immunomodulatory cues, over single-target approaches. Our findings establish early microglial programs as a paradigm for designing new regenerative therapies for CNS disorders.

## Introduction

Neurodegenerative diseases such as multiple sclerosis (MS) and age-related dementia pose significant clinical challenges due to their progressive nature and the current absence of effective regenerative therapies. MS is a chronic inflammatory disorder characterized by demyelination, axonal damage, and progressive neurodegeneration [1]. The disease pathology is further complicated by infiltration of peripheral immune cells into the central nervous system (CNS), exacerbating neurological impairment [2]. Existing therapeutic strategies primarily focus on immunomodulation [3], but they fail to promote remyelination and neuroregeneration [4, 5]. Cognitive decline and memory impairment, common features of normal aging as well as MS, profoundly impact patients’ quality of life [6]. The unmet clinical needs in these and other neurodegenerative conditions underscore the urgency of developing interventions capable of promoting regeneration, cognitive restoration, and functional recovery across diverse CNS disorders.

Microglia, the resident immune cells of the CNS, are crucial in maintaining homeostasis. In the adult brain, they preserve myelin integrity [7], support cognitive functions, mediate immune surveillance, synaptic pruning, and responses to injury [8–11]. Their remarkable plasticity and ability to respond dynamically to diverse environmental cues [12–14] are particularly important in the context of neuroinflammatory and neurodegenerative conditions. Unlike other immune cells, microglia are long-lived and self-renewing, maintaining their population through proliferation [15–17]. However, aging impairs several critical microglial functions, potentially exacerbating neurodegeneration in age-related diseases such as in progressive forms of MS (PMS) or mild cognitive impairment or dementia [18–20]. Despite their essential roles in CNS homeostasis, therapeutic strategies targeting microglial regenerative potential remain largely unexplored.

In addition to their homeostatic roles in adulthood, microglia critically shape brain development. Neonatal microglia (NeoMG) actively drive synaptic pruning [21–23], neurogenesis [24], primary myelination [25, 26] and structural integrity of the developing brain [27]. During this period, they transiently express factors necessary for axonal guidance, myelination, astrocyte differentiation, tissue remodelling and angiogenesis [25]. Understanding and leveraging these developmental regenerative capacities of NeoMG could open novel therapeutic avenues for treating CNS pathologies.

Previously, we and others demonstrated that NeoMG exhibit pronounced neurogenic and myelinogenic capabilities [25, 28–30]. In the present study, we sought to determine whether these trophic, pro-regenerative functions decline with age, and whether restoring them could promote brain repair and functional recovery in MS and ageing-related contexts. To address this, we tested the therapeutic potential of NeoMG across three distinct experimental paradigms: (I) EAE, a model of autoimmune, inflammation-driven demyelination that enables assessment of NeoMGs’ ability to modulate active immune responses in an MS-relevant setting; (II) cuprizone (CPZ)-induced demyelination, which enables assessment of NeoMG-mediated remyelination independent of ongoing inflammation; and (III) a model of physiological ageing associated with cognitive decline. Together, these complementary models enabled us to investigate whether NeoMG can restrain autoimmune pathology, stimulate myelin regeneration, and enhance cognitive function within the age-compromised CNS.

Our findings show that NeoMG transplantation ameliorates EAE pathology, reduces leukocyte infiltration, and enhances myelin regeneration in both inflammatory (EAE) and non-inflammatory (cuprizone) models of demyelination. Remarkably, the same intervention in aged mice reversed memory deficits. Comprehensive multi-omic analyses revealed that these regenerative effects are associated with a NeoMGs’ secretome, uniquely enriched in trophic and metabolic factors, pointing to a paracrine mechanism of action. By re-engaging developmental repair programs NeoMG transplantation parallels rejuvenation paradigms such as heterochronic parabiosis [31–33], where young systemic factors reinstate plasticity and cognition in the aged brain. Collectively, these findings highlight the need to move beyond single-molecule strategies and toward multifactorial therapies that integrate trophic, metabolic, and immunomodulatory pathways. Our study thus establishes NeoMG as a blueprint for harnessing early immune programs to design regenerative interventions for demyelinating and age-related neurodegenerative disorders.

## Results

### Transplantation of neonatal microglia ameliorates EAE and induces remyelination in MS models

In the present study, we evaluated the therapeutic potential of NeoMG in promoting remyelination and modulating disease progression in two murine models of MS: EAE and CPZ.

First, we transplanted microglia MACS-sorted from neonatal and adult mice directly to the cerebrospinal fluid (CSF) via intrathecal injections (i.t.) to the cisterna magna of animals exhibiting tail paralysis, the first symptom of EAE, and monitored the disease progression (Fig. 1A-B).

**Fig. 1.**
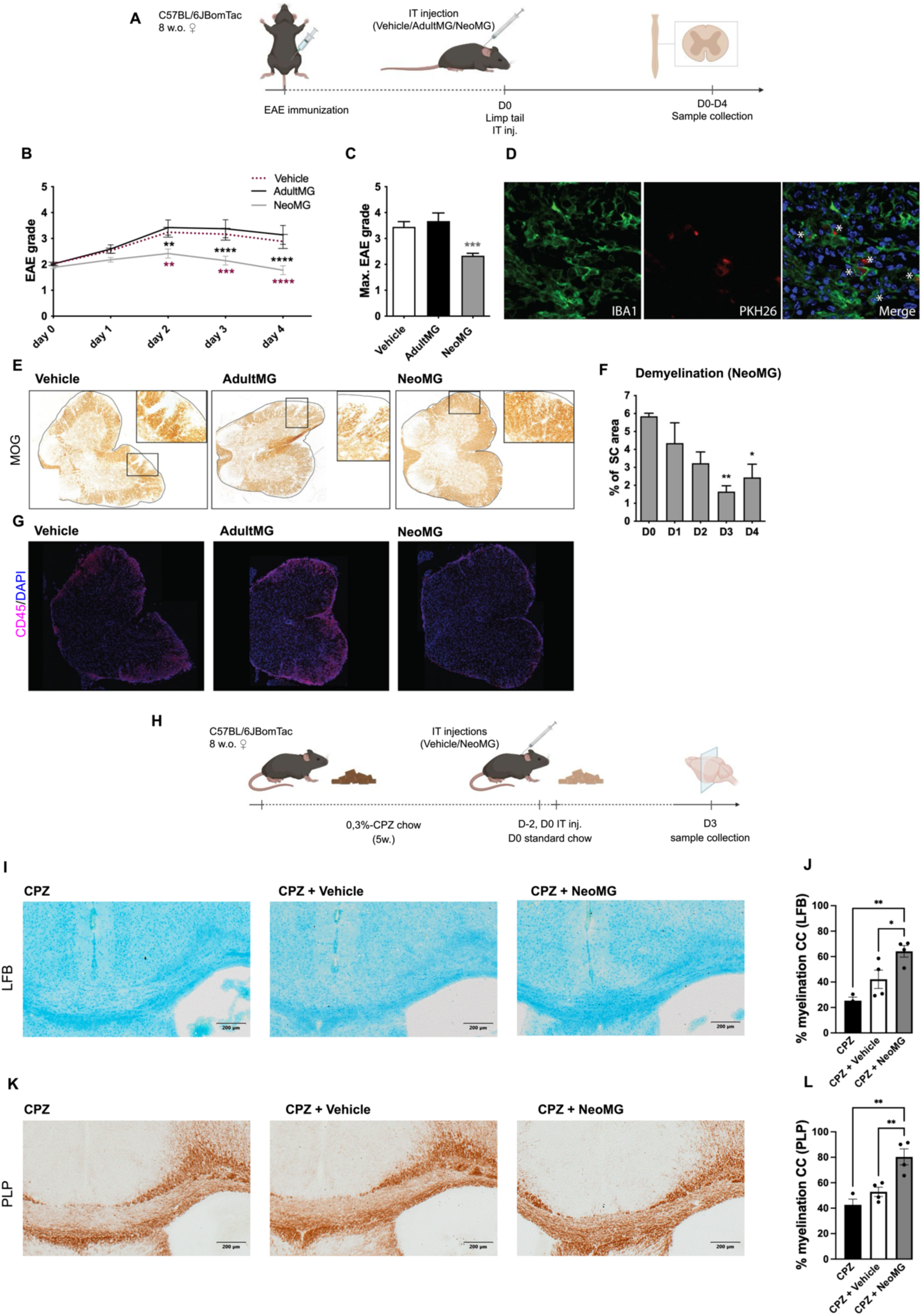
Effects of NeoMG treatment on clinical scores and myelin integrity in EAE and CPZ models. (A) Schematic representation of the experimental setup: EAE-model mice received treatments with NeoMG, AdultMG, or vehicle control (PBS). (B) Impact of NeoMG (gray) AdultMG (black), and vehicle control (red-dotted) treatment on EAE disease progression in 8-10 weeks old C57BL/6 mice (C) The mean maximum grade of EAE symptoms reached 4 days post treatment (D) Representative micrographs of IBA1+ (green) and PKH26-labelled NeoMG (red) intrathecally injected to animals at grade 2 of EAE. White asteriks show IBA1+ cells labelled with PKH26. (E) Myelin loss in spinal cord (SC) of vehicle-, AdultMG- and NeoMG-treated animals 48h after the treatment. Representative SC sections stained for anti-myelin oligodendrocyte glycoprotein (MOG). Myelin is shown as brown-stained white matter area and demyelination is indicated as unstained regions. Magnified areas of demyelination are positioned at the top right corner of each section. (F) Kinetics of demyelination (% of demyelinated SC white matter area) in NeoMG-treated EAE animals 0, 1, 2, 3, and 4 days after the treatment (n=3-4). (G) Infiltration of CD45+ cells in the SC sections of vehicle-, AdultMG- and NeoMG-treated animals. (H) Schematic representation of the experimental setup: CPZ-treated mice received treatments with NeoMG or vehicle (I, K) Representative images of brain sections, showing the extent of demyelination in the CC area, visualised by staining with (I) LFB or (K) anti-PLP. (J, L) Evaluation of myelination by analysis of (J) LFB and (L) anti-PLP staining intensity (n=2-4). Data are expressed as mean ± SEM; each n represents an individual mouse. P-values were determined by two-way ANOVA with Tukey’s test (B, F), Kruskal-Wallis with Dunn’s test (C) or one-way ANOVA with Tukey’s test (J, L). Asterisks indicate significant differences (* P< 0.05; ** P< 0.01; *** P< 0.001; **** P < 0.0001).

Transplantation of adult microglia (AdultMG) did not alter the course of EAE (Fig. 1B); however, NeoMG transplantation significantly halted disease progression and in many cases led to full clinical recovery, manifesting with restored tail mobility. Unlike vehicle-treated controls and AdultMG recipients, mice receiving NeoMG did not progress to more severe EAE stages and did not reach the ethical endpoint (Figure 1B-C). Importantly, transplanted PKH26-labelled NeoMG were detectable by immunostaining within spinal cord lesions for at least 4 days post-transplantation (Fig. 1D). Histological assessments revealed that spinal cords from mice treated with NeoMG showed less severe demyelination (Fig. 1E-F) and reduced leukocyte infiltration (CD45-high) (Fig. 1G) compared to those treated with vehicle or AdultMG. Importantly, our observations revealed a time-dependent reduction of the demyelination area following transplantation of NeoMG (Fig. 1F), suggesting an induction of active remyelination rather than mere suppression of ongoing demyelination.

To further investigate the remyelinating properties of NeoMG, we used a CPZ-induced demyelination model [34]. Mice were fed with chow containing 0.3% of CPZ for five weeks to induce demyelination in the corpus callosum (CC). The treatment resulted in approximately 80% of demyelination (measured by lack of Luxol Fast Blue (LFB) staining in CC) compared to unmanipulated controls (Fig. 1J). Two days prior and on the day of CPZ cessation, we transplanted NeoMG to assess their ability to stimulate naturally occurring remyelination (Fig. 1H). The treatment with NeoMG significantly enhanced remyelination in the CC, as evidenced by increased LFB and Proteolipid Protein (PLP) staining intensities (Fig. 1I-L), confirming the pro-remyelinative properties of NeoMG.

### Transplanted neonatal microglia home to the meninges and improve memory in cognitively impaired animals

Building on the demonstration of the neurosupportive profile and remyelination capabilities of NeoMG, we investigated their potential to reverse cognitive impairments associated with ageing. Utilising 18-22-month-old C57BL/6 mice that naturally exhibit cognitive deficits, we assessed the impact of transplantation of NeoMG on spatial and recognition memory, locomotor activity, and anxiety [35–37] (Fig. 2A).

**Fig. 2.**
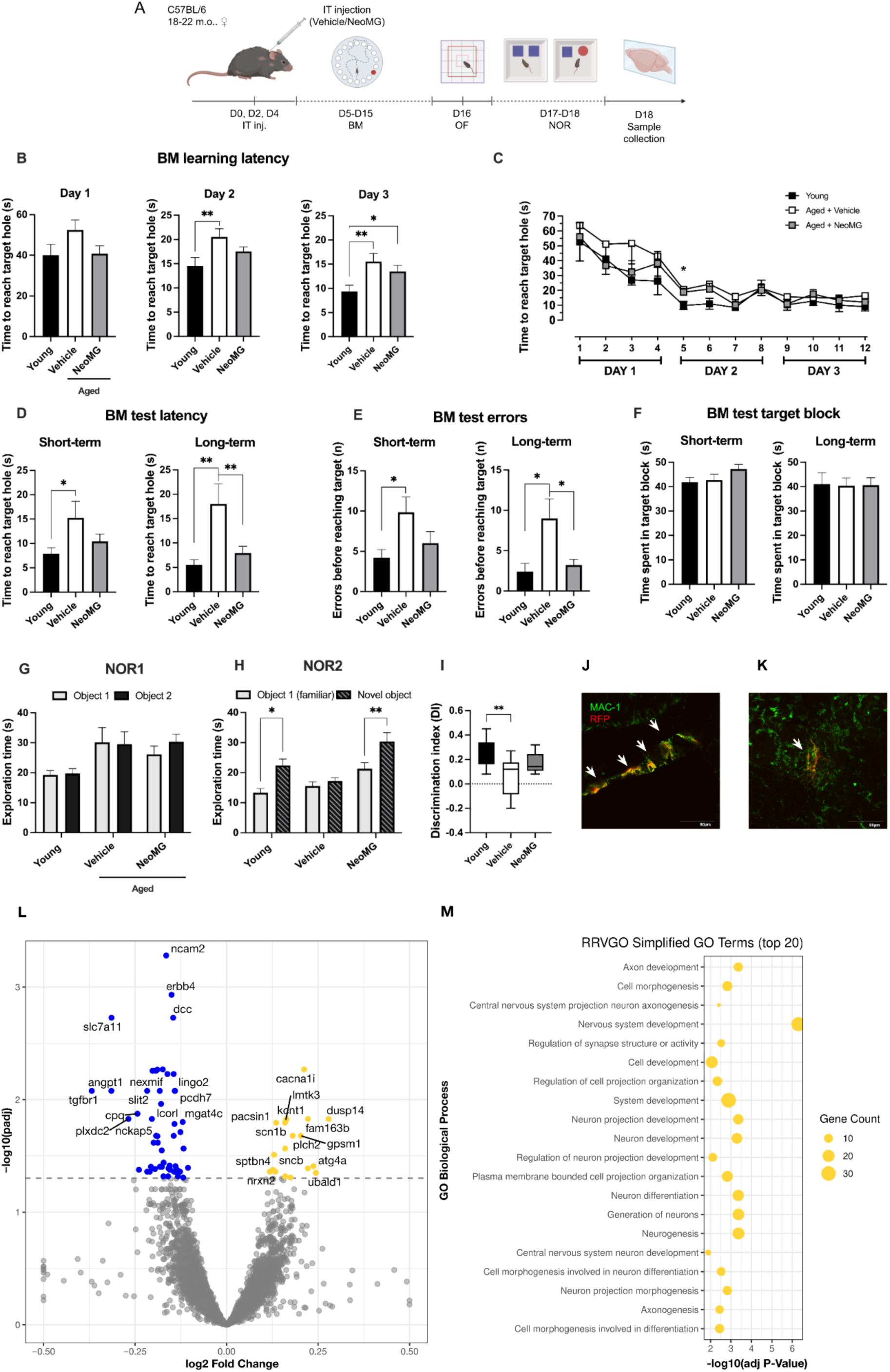
Performance of aged C57BL/6 mice injected with NeoMG vs. PBS-control (vehicle) and young C57BL/6 mice in cognitive behavioural test. (A) Schematic representation of the experimental setup: Intrathecal injections of NeoMG or vehicle to CSF of aged C57BL/6J mice, followed by behavioural testing. (B) Average latency to reach the target hole in the training session (Day 1-3) of the BM task (C) Average latency to reach the target hole during each trial of the training session (Day 1-3) of the BM task. (D) Average latency to reach the target hole during probe trials (Day 4 and 11) of the BM task. (E) Primary error nose pokes done before reaching target in probe trials of the BM task. (F) Average time spent in the maze quarter with the target hole (target block) of the BM task. (G-H) Comparison of the time spent by groups with object 1 (familiar) and 2 (novel) separately, during (G) training and (H) testing phases of NOR task. (I) Discrimination index in the NOR2 (test phase). (J-K) Representative confocal photomicrographs showing MAC-1^+^ (green) and RFP^+^ (red) staining in the meninges (J) and the cerebellum (K) of the RFP^+^NeoMG-treated aged C57BL/6J mice, 1day post-intrathecal injection; scale bars 80 µm. (L) Volcano plot of log₂ fold changes in gene expression between NeoMG-(yellow) and vehicle-treated (blue) C57BL/6J mice. (M) Gene Ontology (GO) enrichment analysis of the significantly differentially expressed genes identified in (L). Data shown as mean ± SEM; each n represents an individual mouse; P-values were determined by one-way ANOVA with Tukey’s test or Kruskal-Wallis with Dunn’s test (B, D-E, H, I) and two-way ANOVA with Tukey’s test (C, F-G). Asterisks indicate significant differences (* p<0.05; ** p<0.01).

Spatial memory was evaluated using the Barnes maze test (BM), where primary latency and errors were measured. As expected, aged mice exhibited significantly impaired learning and short- and long-term memory in comparison to young mice, as previously demonstrated [38–40]. While NeoMG transplantation did not affect learning (Fig. 2B-C) or the total time spent in the target block area (Fig. 2F), shorter latency (Fig. 2D) and fewer errors (Fig. 2E) in short- and long-term memory tests were observed, indicating significantly improved memory in treated aged animals. This performance was comparable to that of young animals, suggesting that NeoMG transplantation led to the restoration of memory in aged animals.

Recognition memory was assessed using a novel object recognition (NOR) task. During the learning phase, on day 1 (NOR1) mice explored two identical objects for 10 min. As expected, no preference for either object was observed in any of the groups on day 1 (Fig. 2G). In the recognition memory test (NOR2) performed 24h post learning, aged mice showed no preference for the novel object and had decreased exploration time, indicating impaired recognition memory (Fig. 2H). In contrast, NeoMG-treated aged animals displayed increased exploration of the novel object, similar to young mice (Fig. 2H). This was supported by a higher discrimination index (DI) in the NeoMG-treated aged mice compared to controls, with no significant difference between young and NeoMG-treated aged mice (Fig. 2I). These results demonstrate that NeoMG treatment reversed recognition memory deficits in aged mice.

Locomotor functions and anxiety were evaluated using the open field test (OF). No significant differences were found between the NeoMG-treated animals and the control group, indicating the treatment does not affect anxiety-related behaviour or locomotor activity (data not shown). Therefore, the observed protective effect of the treatment was specifically related to improvement of memory in aged animals.

Subsequently, we investigated localization of the intrathecally transferred NeoMG within aged brain tissue. To easily discriminate transferred cells, we used CAG::mRFP1 mice (in which all the cells express monomeric red fluorescent protein-1 (mRFP1)) as NeoMG donors. Brain sections were immunostained for macrophage-1 antigen (MAC1) and red fluorescent protein (RFP) to visualise MAC1^+^ RFP^−^ resident microglia and MAC1^+^ RFP^+^ transferred NeoMG.

Confocal microscopy analysis one day after intrathecal injection revealed that transplanted NeoMG cells localised primarily within the meninges and, to a lesser extent, in the cerebellum (Fig. 2J-K), in contrast to EAE where transplanted microglia are found in lesions in the CNS parenchyma.

We next examined the transcriptional changes induced by NeoMG transplantation in the aged brain. Differential gene expression analysis revealed a distinct set of transcripts significantly altered in NeoMG-treated versus PBS-treated (vehicle) aged C57BL/6J mice (Fig. 2L). Gene Ontology (GO) enrichment analysis of the differentially expressed genes highlighted functional categories related to CNS development, neuronogenesis, axonogenesis, cell differentiation and synaptic remodelling, indicating a coordinated transcriptional response following treatment (Fig. 2M).

### Secreted factors are main drivers of the NeoMG-induced therapeutic effects

Given the primary localization of the transplanted NeoMG in the meninges with simultaneous effect on the memory outcomes, it is plausible that secreted factors rather than cell-cell interaction mediate these neuroprotective effects. To test this hypothesis, we administered NeoMG-conditioned media (NeoMG-CM) or control media into the CSF of aged mice and performed behavioural analysis (Fig. 3A).

**Fig. 3.**
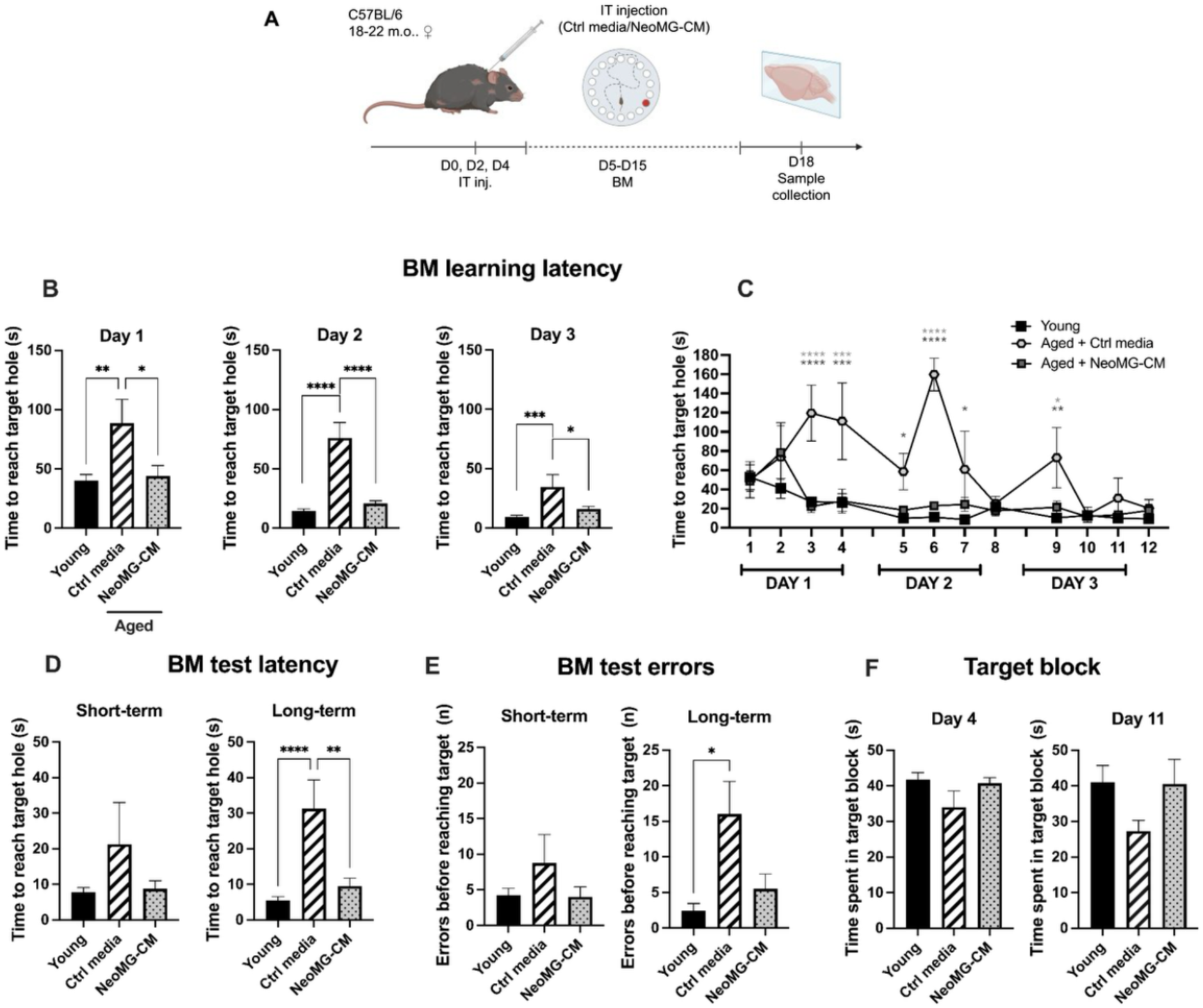
Performance of aged C57BL/6 mice treated with NeoMG-CM vs. control media compared to young C57BL/6J mice in the BM test. (A) Schematic representation of the experimental setup: Intrathecal injections of NeoMG-CM or ctrl media to CSF of aged C57BL/6 mice, followed by behavioural testing using the BM. (B) Average latency to reach the target hole in the training session (Day 1-3). (C) Average latency to reach the target hole in each trial of the training session (Day 1-3). (D) Average latency to reach the target hole in probe trials (Day 4 and 11) of the BM task. (E) Primary error nose pokes done before reaching target in probe trials. (F) Average time spent in the maze quarter with the target hole (target block). Data shown as mean ± SEM; each n represents an individual mouse; P-values were determined by one-way ANOVA with Tukey’s test (B, D-F) or two-way ANOVA with Tukey’s test (C) (* P< 0.05; ** P< 0.01; *** P< 0.001; **** P < 0.0001).

The cognition of the aged animals treated with NeoMG-CM was evaluated using the BM test assessing primary latency and errors. Surprisingly, in contrast to cell transplantation, aged animals receiving NeoMG-CM treatment exhibited improved learning skills, as indicated by shorter latency in identifying the target during learning sessions (Fig. 3B-C). The learning skills of NeoMG-treated animals were comparable to the young animals. Similar to NeoMG transplantation, treatment with NeoMG-CM had a positive effect on memory. Although no significant differences were found in primary latency, errors or time in the target block in a short-term memory test, NeoMG-CM significantly improved long-term memory, with reduced primary latency (Fig. 3D), fewer errors (Fig. 3E), and a trend toward more time spent in the target block (Fig. 3F).

Subsequently, we aimed to determine whether the remyelinating effects of NeoMG are also mediated by their secretome. To test that, we employed the CPZ-induced demyelination model to induce demyelination in the CC during a 5-weeks CPZ treatment. NeoMG-CM was administered i.t. on days –4, –2 and 0 relative to CPZ withdrawal (Fig. 4A). After resumption of normal chow, a 3-day window of spontaneous remyelination was initiated. Remarkably, treatment with NeoMG-CM significantly enhanced remyelination in the CC, as evidenced by increased LFB staining intensity, following this 3-day period (Fig. 4B-C). This finding highlights the potent pro-remyelinating effect of factors secreted by NeoMG.

**Fig. 4.**
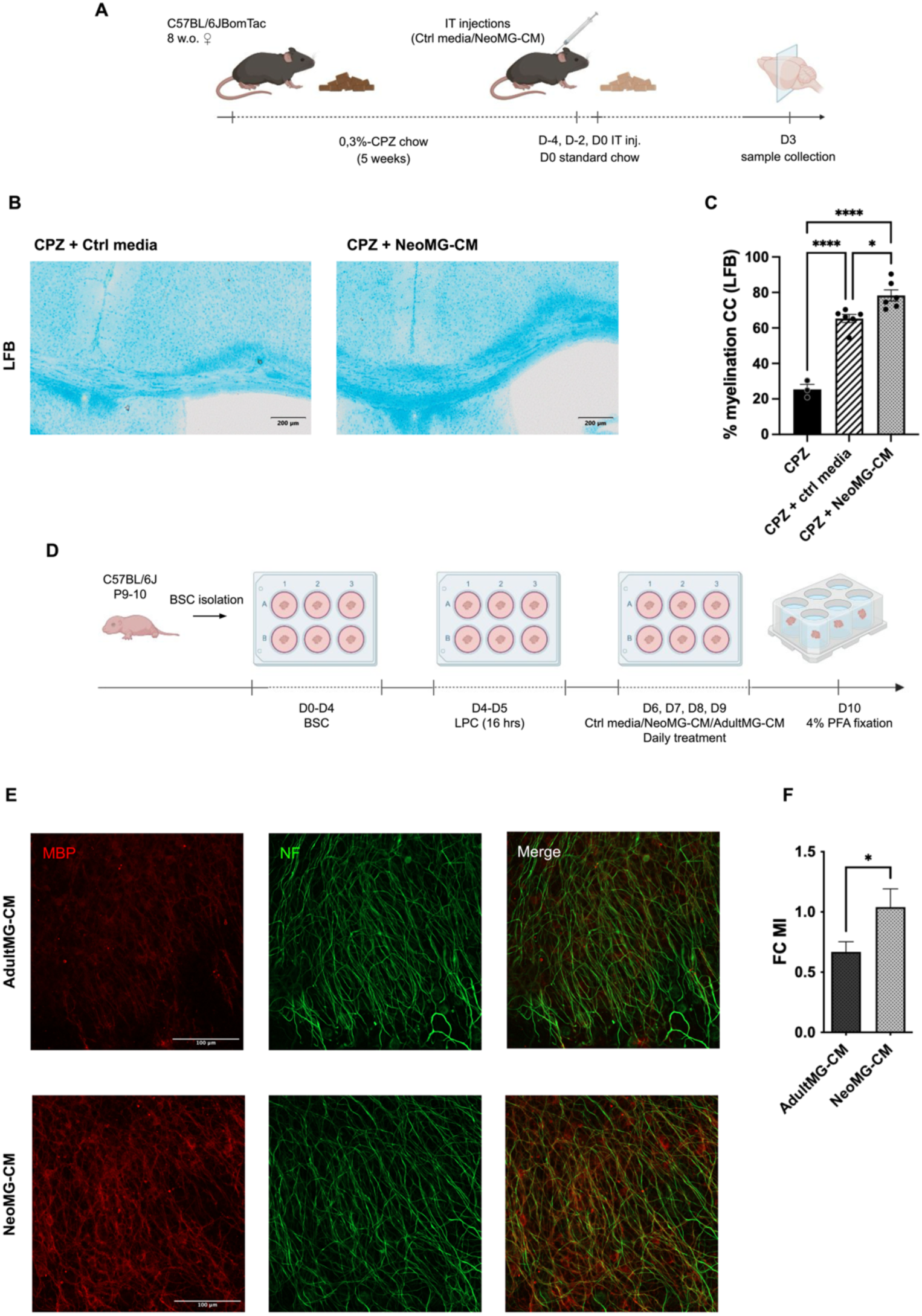
Effects of NeoMG-CM treatment on remyelination in CPZ model and BSC. (A) Schematic representation of the experimental setup: Intrathecal injections of NeoMG-CM or control media to CSF of CPZ-treated animals. (B) Extent of demyelination in CC of the LFB-stained brain sections after 5-6 weeks of CPZ intoxication, with control media or NeoMG-CM treatment, and 3 days of remyelination following CPZ withdrawal. (C) Graph bars showing LFB staining intensity of myelination in CC across treated groups (n=3-7). (D) Schematic representation illustrating the isolation and culture of cerebellar brain slices, followed by the LPC-induced demyelination and treatment with NeoMG-CM or AdultMG-CM. (E) Representative immunofluorescence images of cerebellar brain slices stained with anti-myelin basic protein (MBP, red) and anti-neurofilament (NF, green) following treatment with NeoMG-CM or AdultMG-CM. Scale bars 100 µm. (F) Fold change (FC) in myelination index (MI), calculated as the relative number of MBP⁺/NF⁺ axons over total NF⁺ axons in slices treated with NeoMG-CM or AdultMG-CM, normalized to control medium-treated slices; (n =8-10 slices per group, isolated from ≥ 3 pups of mixed sex). Data shown as mean ± SEM; each n represents an individual mouse (C) or cerebellar brain slice (F). P-values were determined by one-way ANOVA with Tukey’s test (C) and unpaired two-tailed t-test (F) (* P< 0.05; ** P< 0.01; *** P< 0.001; **** P< 0.0001).

To test whether the improved remyelination seen with NeoMG-CM treatment in the CPZ model reflects age-specific secreted factors, we compared NeoMG-CM and AdultMG-CM in an ex-vivo cerebellar brain-slice culture (BSC) model. Slices were demyelinated locally with lysolecithin (LPC) [41] and 24h later treated daily with either NeoMG-CM or AdultMG-CM for 4 days (Fig. 4D). NeoMG-CM-treated slices showed higher co-localization of myelin basic protein (MBP) with neurofilament (NF) fibres than slices receiving AdultMG-CM (Fig. 4E). Quantification confirmed a significantly greater myelination index in the NeoMG-CM-treated slices (Fig. 4F), suggesting that NeoMG-CM, but not AdultMG-CM promote remyelination. Overall, the use of CPZ-and LPC-induced demyelination models demonstrated that NeoMG-CM treatment promotes remyelination in both systemic (CPZ) and localized (LPC) contexts [36]. Altogether, our findings show that the beneficial effects of NeoMG-CM transplantation can be attributed to extracellular factors.

### Comparative Proteomic Analysis of Neonatal and Adult Microglia Conditioned Media

To define molecular differences underlying the distinct therapeutic capacity of NeoMG, we performed comparative proteomic profiling of conditioned media (CM) from NeoMG and AdultMG. Principal component analysis revealed clear separation between the two groups, indicating marked divergence in their secretomes (Fig. 5A).

**Fig. 5.**
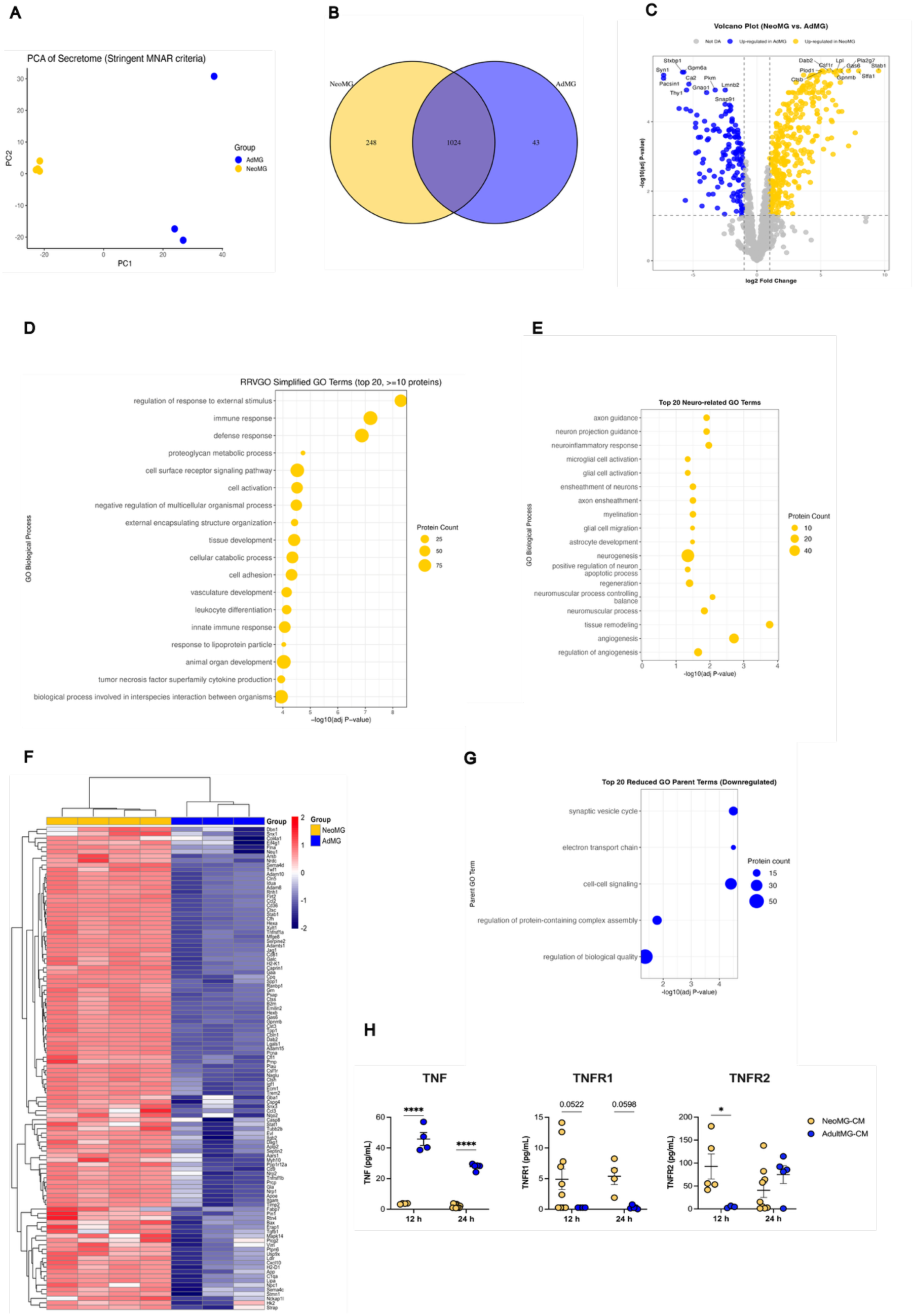
Differential expression of NeoMG-CM and AdultMG-CM proteome. (A) Principal Component Analysis (PCA) of proteomic profile in NeoMG-CM and AdultMG-CM. (B) Venn diagram illustrating the overlap of 1019 proteins, 246 proteins uniquely expressed in NeoMG-CM and 42 proteins uniquely expressed in AdultMG-CM. (B) Volcano plot showing the log2 fold change (x-axis) against the t-test–derived −log10 statistical p-value (y-axis) for all enriched proteins differentially expressed between NeoMG-CM and AdultMG-CM proteome. Selected enriched proteins are labelled. (D) Gene Ontology (GO) enrichment analysis of differentially expressed proteins enriched in NeoMG-CM. (E) Enriched CNS-related GO terms (n=18) among proteins upregulated in NeoMG-CM. Terms are ranked by –log₁₀(adjusted P-value); dot size indicates the number of proteins per term. (F) Heatmap depicting CNS-related differentially expressed proteins in each sample of NeoMG-CM and AdultMG-CM. Protein expression levels per sample are shown relative to the average expression across all samples. The colour key corresponds to row Z-scores, where red indicates higher expression and blue indicates lower expression. (G) Enriched GO terms among proteins upregulated in AdultMG-CM. Categories are ranked by – log₁₀(adjusted P-value); dot size reflects protein count. (H) solTNF, solTNFR1, solTNFR2 concentrations in NeoMG-CM and AdultMG-CM at 12- and 24-h-time points of CM collection. Data represent mean ± SEM; n=3– 9 technical replicates from 1–2 independent biological experiments (each biological replicate pooled from >= 3 animals). Statistical analysis was performed using two-way ANOVA followed by Fisher’s LSD post hoc test; (* P< 0.05, ** P< 0.01, *** P< 0.001, **** P< 0.0001).

Our analysis identified a total of 1,272 proteins in NeoMG-CM and 1,067 proteins in AdultMG-CM. Notably, 358 proteins were significantly upregulated in NeoMG-CM (p ≤ 0.05, Log2FC ≥ 1), among which 248 proteins were uniquely expressed in neonatal samples (Suppl. Table S1). Conversely, AdultMG-CM showed 144 significantly higher expressed proteins, with 43 proteins uniquely expressed (Fig. 5B). Proteins enriched in NeoMG-CM (Fig. 5C) corresponded to 363 distinct Gene Ontology (GO) terms (Suppl. Table S1). This broad spectrum of enriched GO terms underscores the pleiotropic biological functions inherent to NeoMG. Semantic reduction refined these GO terms to 115 representative categories, prominently including immune response, tissue and organ development, vascular development, cell activation, and cytokine production within the tumor necrosis factor (TNF) superfamily (Fig.5D).

Importantly, enriched terms also captured processes directly relevant to CNS repair. Keyword extraction revealed 18 significantly overrepresented brain-related categories, including axon guidance, microglial activation, myelination, astrocyte development, regeneration, tissue remodeling, and angiogenesis (Fig. 5E). Filtering for proteins annotated to these terms identified 109 CNS-relevant candidates among the NeoMG-enriched set (Fig. 5F). In contrast, proteins enriched in AdultMG-CM were associated with only 50 GO terms (Suppl. Table S1), reduced to five functional categories: electron transport chain, synaptic vesicle cycling, cell–cell signalling, regulation of protein complex assembly, and regulation of biological quality (Fig. 5G).

Given the enrichment of TNF-associated pathways in NeoMG-CM, we further quantified soluble TNF (solTNF) and its receptors. While solTNF levels were higher in AdultMG-CM, NeoMG-CM contained significantly greater amounts of soluble TNFR2 and a trend toward increased solTNFR1, particularly at early collection time points (Fig. 5H). This profile is consistent with preferential engagement of protective TNFR2-mediated signalling, while limiting detrimental solTNF/TNFR1 activity.

Taken together, these data reveal that the NeoMGl secretome is distinguished by a broader and more functionally diverse proteomic repertoire than that of adults, enriched for pathways directly linked to CNS development and repair. These findings suggest that the therapeutic potential of NeoMG derives from a complex interplay of neurotrophic, immune-regulatory, and metabolic signals, rather than the action of individual factors.

### Comparative Lipidomic Analysis of Neonatal and Adult Microglia Conditioned Media

As microglial lipid metabolism shifts with age and contributes to neurodegenerative vulnerability [42], we investigated whether lipid composition underlies the enhanced regenerative capacity of NeoMG. Comparative analysis of the cellular and secreted fractions revealed marked differences between NeoMG and AdultMG (Fig. 6A-F). In both populations, diacylglycerol (DG) was the most abundant class. NeoMG contained proportionally more phosphatidylcholine (PC), phosphatidylethanolamine (PE), and sphingomyelin (SM), while AdultMG exhibited higher DG and reduced phospholipid content (Fig. 6A–C). Volcano plot analysis further showed significant enrichment in NeoMG of ether-linked phospholipids (plasmalogens: PE-P, PC-P; ether-linked PE-O, PC-O), lysophospholipids (LPE, LPC), and hexosylceramides (HexCer) (Fig. 6F). These lipids are essential for membrane biosynthesis, myelin integrity, and oxidative stress resistance [43–48], suggesting a metabolic state primed for regeneration. In contrast, AdultMG were enriched in neutral storage and stress-associated lipids DG, Cer, and certain TG and PE-P species (Fig. 6C, F) consistent with a less reparative phenotype.

**Fig. 6.**
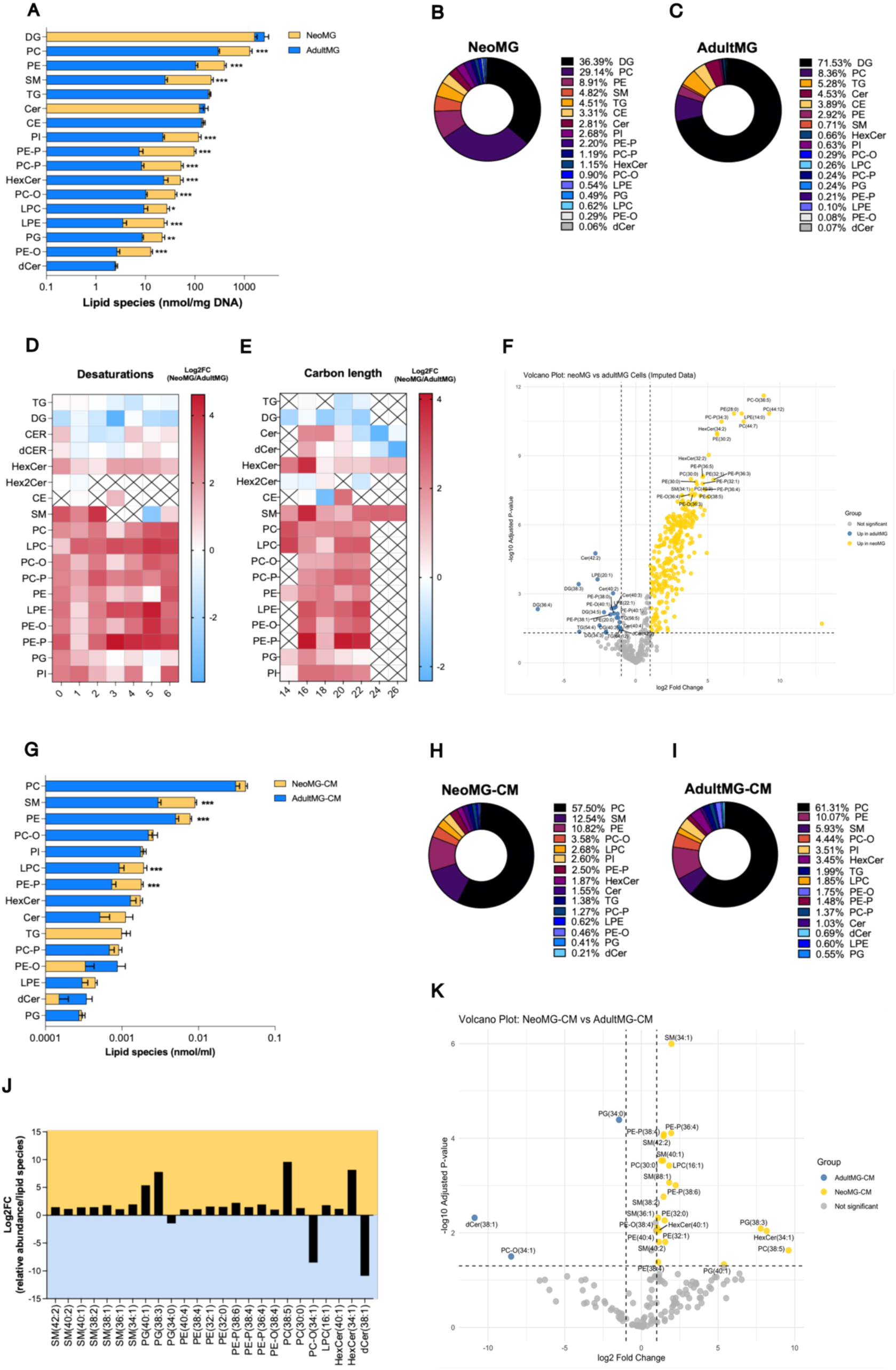
Lipidomic profiling of NeoMG, AdultMG, and their CM. (A) Concentration of major lipid classes in NeoMG and AdultMG, shown as scaled bar graphs. Bars represent mean ± SEM. Statistical significance was assessed using multiple unpaired t-tests followed by Benjamini-Hochberg correction for multiple comparisons (FDR < 1%). Significant differences are indicated as follows: *P< 0.05; ** P< 0.01; *** P< 0.001. (B-C) Proportional distribution of lipid classes in NeoMG (B) and AdultMG (C), expressed as percentage of total lipid abundance. (D-E) Heatmaps of lipid species detected in NeoMG and AdultMG grouped by carbon chain length (B) and degree of saturation. (E) Heatmaps display log₂ fold change (NeoMG/AdultMG) in lipid abundance. Color intensity reflects the magnitude and direction of change, with red indicating increased levels in NeoMG and blue indicating increased levels in AdultMG. (F) Volcano plot analysis of lipid species differentially abundant between NeoMG and AdultMG. The x-axis represents log₂(fold change); the y-axis shows –log₁₀(P-value). Significantly altered lipids (adjusted P < 0.05, Benjamini–Hochberg correction) are highlighted; lipids enriched in AdultMG appear in blue, and those enriched in NeoMG appear in yellow. (G) Concentration of major lipid classes in NeoMG-CM and AdultMG-CM, shown as scaled bar graphs. Bars represent mean ± SEM. Statistical significance was assessed using multiple unpaired t-tests followed by Benjamini-Hochberg correction for multiple comparisons (FDR < 1%). Significant differences are indicated as follows: * P< 0.05; ** P< 0.01; *** P< 0.001. (H-I) Proportional distribution of lipid classes in NeoMG-CM and AdultMG-CM, expressed as percentage of total lipid abundance. (J) Lipid species with significantly different abundance between NeoMG-CM and AdultMG-CM, shown as log₂(fold change) values. (K) Volcano plot analysis of lipid species differentially abundant between NeoMG-CM and AdultM-CM. The x-axis represents log₂(fold change); the y-axis shows –log₁₀(P-value). Significantly altered lipids (adjusted P < 0.05, Benjamini–Hochberg correction) are highlighted (J-K); lipids enriched in AdultMG-CM appear in blue, and those enriched in NeoMG-CM appear in yellow. Lipid class abbreviations: dihydroceramide (dCer), triacylglycerol (TG), sphingomyelin (SM), phosphatidylinositol (PI), phosphatidylglycerol (PG), plasmenyl phosphatidylethanolamine (PE-P), ether-linked phosphatidylethanolamine (PE-O), phosphatidylethanolamine (PE), plasmenyl phosphatidylcholine (PC-P), ether-linked phosphatidylcholine (PC-O), phosphatidylcholine (PC), lysophosphatidylethanolamine (LPE), lysophosphatidylcholine (LPC), hexosylceramide (HexCer), dihexosylceramide (Hex2Cer), diacylglycerol (DG), ceramide (Cer), and cholesterol esters (CE).

Conditioned media analysis revealed equally striking differences in secreted lipid profiles (Fig. 6G-K). NeoMG-CM was significantly enriched in SM, PE, plasmalogens, and ether-linked phospholipids, whereas AdultMG-CM contained fewer membrane-associated species and more neutral lipids. These membrane-associated lipid classes are well-established components of extracellular vesicles (EVs) (e.g., SM, plasmalogens, cholesterol) [49–51], raising the possibility that EVs may contribute to the regenerative activity of NeoMG secretome. Indeed, NeoMG-CM exhibited robust upregulation of canonical EV markers, including tetraspanins (CD9, CD63, CD81), ESCRT-associated proteins (TSG101, SDCBP), and RNA-binding proteins implicated in EV cargo loading (YBX1), (Suppl.Fig. S1). These proteomic findings align with the EV-like lipidomic signature, together providing convergent evidence that the NeoMG secretome contains abundant EVs.

Collectively, these results demonstrate that NeoMG differ profoundly from AdultMG in both lipid and protein composition, adopting a metabolic and secretory state that favours membrane biosynthesis and EV-mediated communication. The enrichment of EV-associated lipids and proteins in NeoMG-CM suggests that EVs may represent a mechanism through which NeoMG deliver regenerative cues to the aged or demyelinated CNS.

### Neonatal Microglia Transplantation Modulates Ageing-Associated Transcriptomic Signatures via Secretome-Enriched Pathways

To further explore how NeoMG transplantation modulates age-associated transcriptional programs, we integrated transcriptomic signatures from aged brain tissue with proteomic profiles of the NeoMG secretome. GO enrichment analysis revealed strong convergence between pathways altered in the aged CNS and those represented in NeoMG-CM. GO enrichment analysis revealed strong convergence between pathways altered in the aged CNS and those represented in NeoMG-CM (Fig. 7A). Genes and proteins intersecting across datasets clustered into functionally coherent modules, including axonogenesis and neurite extension, glial differentiation, SMAD/TGF-β signalling, and extracellular matrix remodelling. Notably, several of these categories are directly linked to developmental myelination and tissue regeneration. Other clusters highlighted regulation of oxidative stress, cell survival, and immune response pathways, indicating that NeoMG-derived factors may act broadly to reset both trophic and inflammatory networks in the aged brain.

**Figure 7.**
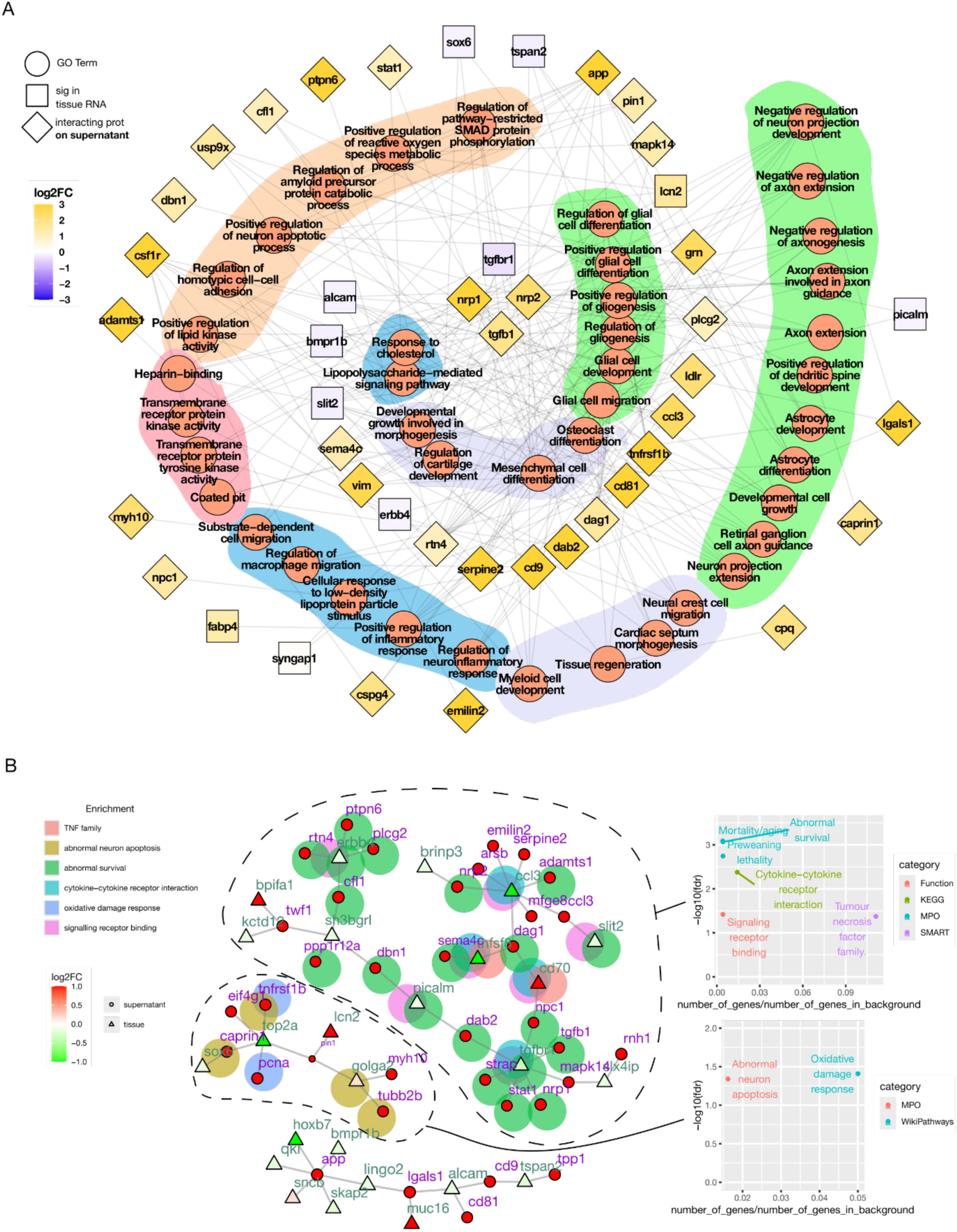
Convergent transcriptomic and proteomic signatures in aged brain tissue and microglial secretome, with functional clustering and network enrichment analysis. (A) GO enrichment analysis of genes that are significantly differentially expressed in aged brain tissue and at the protein level in NeoMG-CM, indicating converging pathways between the two datasets. Background shades indicate groups of GO terms by common ontology ancestry: in orange specific signal transduction, in pink kinase-mediated cellular homeostasis regulation, in green large phenotypic changes, in violet tissue regeneration, in blue specific neuroinflammatory terms. (B) Interactome network connecting these two datasets. Triangles represent genes expressed in the aged brain after NeoMG transplantation and circles represent those from the NeoMG secretome. Only clusters with more than two nodes are shown. Enrichment analysis was performed for the entire network (bottom panel) and for individual clusters (side panels). Highlighted points within the network indicate clusters with significant GO term enrichment.

An interactome network connecting genes differentially expressed in the aged brain after NeoMG transplantation and proteins present in the NeoMG-CM revealed multiple functionally enriched clusters (Fig. 7B). Among these, three prominent clusters were identified: one centred around TGF-β signalling components, a second enriched for genes involved in abnormal neuronal apoptosis and survival, and a third organized around TNF superfamily (*Tnfsf8*, *Cd70*), regulators of neuronal apoptosis (*Sox6*, *Top2a*, *Golga2*, *Tubb2b*), and survival-related genes (*Erbb4*, *Rtn4*, *Plcg2*, *Cfl1*). Additional clusters were associated with cytokine–cytokine receptor interactions (*Sema4c*, *Ccl3*, *Tgfbr1*, *Cd70*), oxidative damage response (*Tnfrsf1b*, *Pcna*), and signalling receptor binding (*Picalm*, *Tnfsf8*, *Ccl3*, *Cd70*, *Slit2*) (Fig. 7B, side panels).

Together, these convergent modules demonstrate that NeoMG factors engage a multifactorial repair program, simultaneously attenuating apoptotic and inflammatory pathways while reactivating developmental and regenerative processes in the aged CNS.

### NeoMG transplantation counteracts ageing-associated transcriptional changes

To determine how NeoMG transplantation modulates age-associated transcriptional programs, we compared gene expression in NeoMG-versus vehicle-treated aged mice alongside age-related changes between 18- and 4-month-old MODEL-AD animals. Scatterplot analysis (Fig. 8A) revealed that many transcripts upregulated during ageing were suppressed following NeoMG treatment (quadrant 4), whereas a smaller subset of age-suppressed transcripts was re-induced (quadrant 1) (Fig. 8B,C). Genes were color-coded by relative protein abundance in NeoMG-CM versus AdultMG-CM, showing that secretome-enriched proteins were disproportionately represented among these compensatory changes, linking secreted factors to transcriptomic shifts in treated animals.

**Figure 8.**
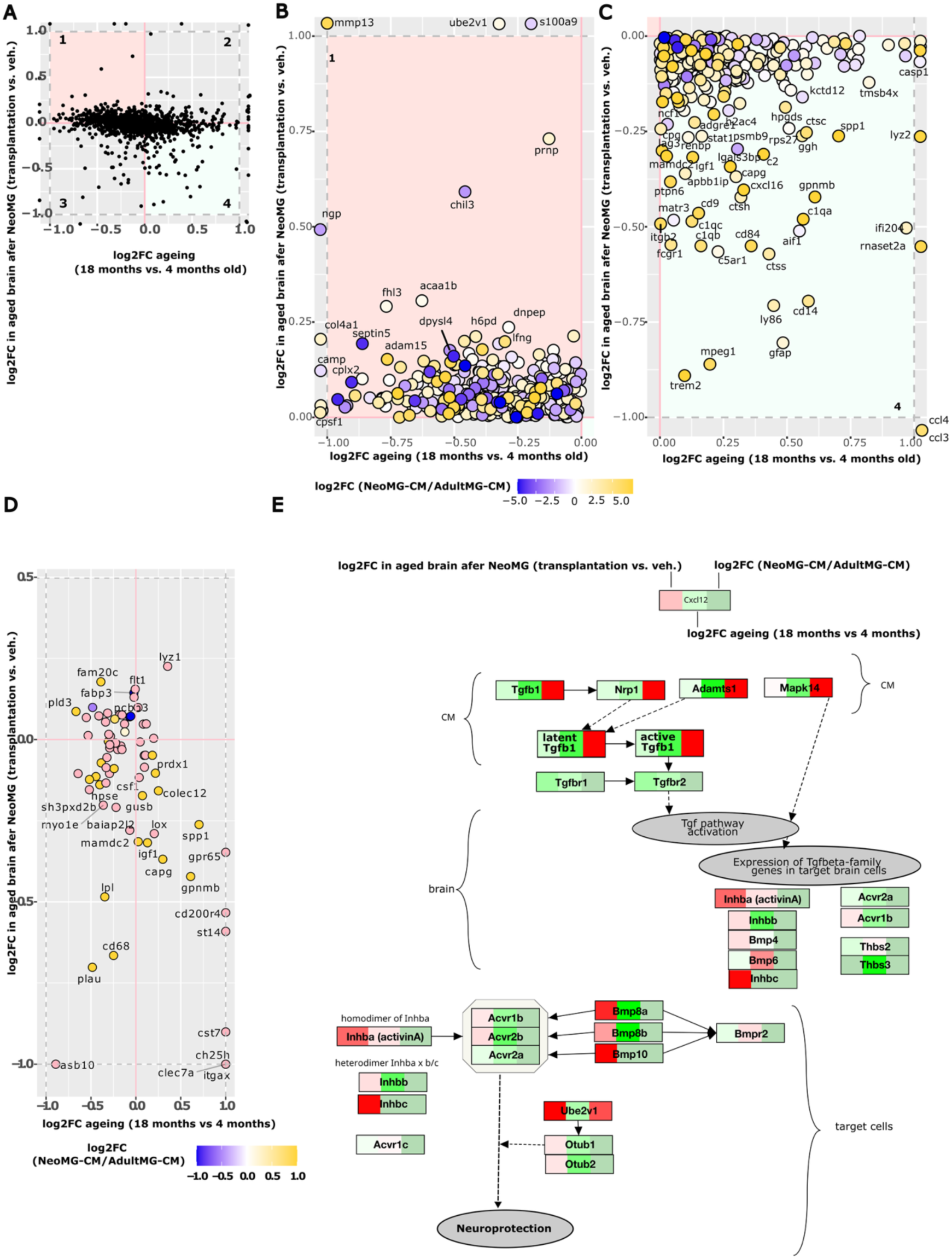
Comparison of gene expression changes in ageing and in the aged brain following NeoMG transplantation, with proteomic annotation from NeoMG-CM versus AdultMG-CM. (A) Scatter plot showing log₂ fold changes in gene expression between NeoMG- and vehicle-treated aged mice (y-axis) and between 18-month-old and 4-month-old mice (x-axis). Genes in the upper left quadrant (1) are downregulated with ageing and upregulated following NeoMG transplantation, while those in the lower right quadrant (4) show the opposite pattern-upregulated with ageing and downregulated following treatment. Each point represents a gene and is coloured according to its relative protein abundance in the NeoMG-CM versus AdultMG-CM, with yellow indicating higher abundance in NeoMG-CM and blue indicating lower abundance. (B, C) Magnified views of the upper left (B) and lower right (C) quadrants of the scatter plot, highlighting genes with inverse expression patterns in ageing and following NeoMG transplantation. (D) Relationship between gene expression changes in ageing and following NeoMG transplantation, restricted to CD11c+ microglia /DAM–associated genes [52]. Genes not detected in the conditioned medium (CM) are indicated in pink. (E) Schematic showing relationship between gene expression changes in ageing and following NeoMG transplantation within Tgf-β/ActivinA pathway [29, 58, 59]. Log₂ fold changes are represented as heatmap colours, with red indicating activation and green indicating inhibition. The left bar shows Log₂FC in the aged brain following NeoMG transplantation, the middle bar shows changes associated with ageing, and the right bar shows differences in the CM.

### Three gene-expression patterns emerged from this analysis

First, genes upregulated in NeoMG-CM and with age, but suppressed after treatment, were enriched for the CD11c⁺/disease-associated microglia (DAM) signature [52], including *Igf1, Spp1, Gpnmb, Itgax, Clec7a*, upregulation of *Aif1*, and *Trem2* (Fig. 8C, D). We also observed a decrease of markers of astrocyte reactivity (*Gfap*) and the complement cascade (*C1qa/b/c, C2, C5ar1*) (Fig. 8C). Several of the DAM-associated genes, which increase with ageing and neurodegeneration, were reduced following NeoMG treatment despite being present in the NeoMG secretome (Fig. 8C, D), suggesting that the DAM-like program diminishes as the tissue transitions toward repair. Second, a subset of genes downregulated in NeoMG-CM but increased after transplantation, including *S100a9, Chil3 (Ym1),* and *Ngp*, are typically enriched in the young brain (Fig. 8B), suggesting activation of endogenous protective immune responses and repair programs [53, 54].

Third, genes upregulated in both NeoMG-CM and aged brains after transplantation included *Mmp13, Ube2v1,* and *Prnp* (Fig. 8B), which regulate extracellular matrix remodelling, ubiquitination in signaling, and neuronal survival, and are normally abundant in the young brain [55–57].

Further pathway analysis revealed prominent representation of the TGF-β/Activin signalling axis (Fig. 8E). Pathway activators present in NeoMG-CM (*TGF-β, NRP1, ADAMTS1, MAPK14*), together with ligands (*Inhba, Bmp8, Bmp10*), receptors (*Acvr1b, Acvr2b*), and regulators (*Ube2v1, Otub1/2*), all showed inverse age regulation and restoration after transplantation. This coordinated re-engagement of Activin/TGF-β signalling and suppression of *Bmpr2* is predicted to drive neurogenesis via stabilization of SMAD2/3 phosphorylation [29, 58].

Together, these findings indicate that NeoMG transplantation exerts multimodal effect on the aged CNS: suppression of the age-induced DAM/CD11c⁺ program, activation of an Activin/TGF-β-driven developmental repair pathway, regulation of astrocyte activation and reduction of complement cascade. This combination likely underlies the observed shift from a pro-inflammatory to a regenerative microenvironment that supports cognitive and functional recovery.

## Discussion

Our results demonstrate that NeoMG and their secretome restore functional homeostasis in aged, demyelinated, and inflamed CNS. They rebalance inflammatory and metabolic networks and enable remyelination, physical and behavioural recovery. This extends prior work on microglial heterogeneity [25, 60] and depletion-repopulation strategies [61, 62] establishing the neonatal repair program as a potent, yet unexplored, therapeutic axis. Notably, i.t administration of NeoMG-CM was sufficient to restore recognition and spatial memory in aged mice, despite limited parenchymal engraftment, underscoring the power of paracrine signalling. In contrast, in EAE, NeoMG infiltrated demyelinated lesions, suggesting a context-dependent dual mode of action: secreted factors in the intact brain, and lesion-directed migration under inflammatory conditions.

Multi-omics analysis identified a distinct secretory program in NeoMG, enriched for neurotrophic and myelination factors (IGF1), axon-guidance molecules (semaphorins), and factors involved in ECM remodelling (MMP13), alongside lipids essential for membrane biogenesis (PC, PE, SM, plasmalogens). Integration of proteomic and lipidomic data suggests a coordinated trophic and metabolic mechanism: soluble factors that promote oligodendrocyte maturation and stimulate myelination, together with lipid mediators and transporters (e.g. APOE, LPL, FABP7, DAB2) that support substrate redistribution and myelin synthesis. This is consistent with evidence for microglial regulation of oligodendrocyte lipid profile [7], oligodendrocyte differentiation and myelination [29, 30]. AdultMG, by contrast, show a lipid-storage phenotype enriched in ceramides and diacylglycerols, consistent with a stress-associated rather than regenerative state. These findings highlight lipid homeostasis and intercellular lipid supply as critical determinants of microglial reparative capacity, which diminishes with age. Importantly, restoration of oligodendrocyte homeostasis via young CSF infusion has been linked to functional brain rejuvenation [63], suggesting that the ability of NeoMG to normalize oligodendrocyte function may represent a convergent mechanism underlying their broad regenerative effects.

The composition of NeoMG-CM suggests the involvement of an EV component. Canonical EV proteins (CD9, CD81) and EV-enriched lipids (SM, plasmalogens) [49, 50] were over-represented, raising the possibility that vesicles mediate concentrated transfer of lipids and signalling molecules. Notably, some of these lipid classes overlap with those enriched in NeoMG-CM and are core components of compact myelin (reviewed in [51]). Prior in vitro studies have shown that EVs from IL-4-polarized microglia or macrophages promote oligodendrocyte precursor recruitment, differentiation and myelination [64, 65]. The overlap between known EV cargo and lipids enriched in NeoMG-CM points to EV-associated signalling as a plausible vehicle for regenerative cues, although future work will need to define vesicle subclasses, cargo and causal mechanisms.

Transcriptomic profiling of treated aged brains revealed suppression of inflammatory mediators (*Trem2, Ccl3, Gfap, Aif1, Cd14*) and induction of tissue-repair genes (*Mmp13, Chil3, Golga2*), as well as downregulation of *SOX6*, a transcriptional repressor of oligodendrocyte differentiation [66], associated with failed remyelination in MS lesions and aged white matter [67]. These changes are consistent with re-engagement of developmental repair pathways [68, 69]. A large proportion of suppressed transcripts overlapped with signature of the CD11c⁺/DAM population, which expands during ageing and neurodegeneration [52]. Their reduction following NeoMG treatment may reflect resolution of the DAM-like state as tissue repair progresses. This is consistent with observations in cuprizone and EAE, where CD11c⁺ microglia peak during active pathology, and decline as repair is established [52, 70–72]. While this association suggests that decreased DAM signatures accompany regenerative outcomes, it remains uncertain whether these changes represent a causal mechanism or a read-out of successful regeneration.

Pathway analyses further implicated the Activin/TGF-β axis, with signalling biased toward neurogenesis and improvements in synaptic plasticity [73–75], and/or pro-remyelinating SMAD2/3 cascades [29, 58]. In treated aged brains, NeoMG transplantation led to upregulation of key pathway components, (*Inhba, Bmp8/10, Acvr1b, Acvr2b),* indicating induction of Activin/TGF-β signalling within the treated brain. Consistent with this, NeoMG-CM contained TGF-β together with its modulators NRP1/2 and DAB2, which act to enhance receptor activation and suppress Toll-like receptor signalling [76–79]. In parallel, NeoMG-CM delivered soluble TNFR2, that is known to selectively attenuate detrimental solTNF signalling, while preserving protective TNFR2 pathways [80–82]. The observed suppression of the degenerative CCL3 inflammatory axis (elevated in activated microglia in ageing [83]), further emphasizes the multifaceted anti-inflammatory properties of NeoMG-CM [84]. Suppression of this pathway has also been shown to have a stimulating effect on remyelination, following deletion of TNFR1 or inhibition of solTNF [85]. Of note, both soluble TNF and CCL3 drive leukocyte recruitment to the CNS, so their coordinated inhibition offers a possible explanation for the reduced immune-cell infiltration observed in EAE. Together, these convergent shifts suggest that NeoMG act by both resolving injury-associated inflammation and activating pro-regenerative signalling cascades.

Altogether, our findings establish NeoMG as potent drivers of CNS repair, but translation will require strategies that capture their activity in clinically viable forms. The identification of key pathways and factors offers an opportunity to design therapeutics that mimic their regenerative profile. Rather than relying on a single target, our data highlight the importance of multifactorial approaches that combine trophic support, lipid redistribution, and immunomodulation. Although the precise composition and impact of NeoMG-derived signals will likely vary across pathological contexts, our results provide a framework for systematically dissecting these mechanisms. By re-engaging developmental repair programs in the adult CNS, NeoMG create an environment supportive for remyelination, cognitive and functional recovery. These findings underscore the therapeutic promise of harnessing early immune programs to develop regenerative strategies for demyelinating and age-related neurodegenerative disorders.

## Methods

### 1. Animals

C57BL/6JBomTac female mice, aged 7-9 weeks, were purchased from Taconic Europe A/S (Denmark). Aged C57BL/6JRj female mice were purchased from Janvier labs (France). 9 weeks old CAG::mRFP1 or C57BL/6J female and male mice were purchased from The Jackson Laboratory (United States) and maintained as a breeding colony to obtain CAG::mRFP1 or C57BL/6J neonates, respectively. CAG::mRFP1, C57BL/6JBomTac neonatal mice (P3-5) and C57BL/6J pups (P9-10) used for experiments were of mixed sex. Animals were maintained and aged in the Biomedical Laboratory at the University of Southern Denmark, at 25°C with 55% humidity on a 12-h light-dark cycle and given free access to food and drinking water. All animal experiments were approved by the Danish Animal Inspectorate (licence number TOW-2020-15-0201-00651) and Ethical Committee for Animal Experiments of Hasselt University (202408K; *ex vivo* use).

### 2. EAE model

Eight- to ten-week-old C57BL/6JBomTac female mice were immunized by subcutaneously injecting 100 μl of an emulsion containing 300 μg of myelin oligodendrocyte glycoprotein (MOG)p35–55 (TAG Copenhagen A/S, Frederiksberg, Denmark) in incomplete Freund’s adjuvant (DIFCO, Albertslund, Denmark) supplemented with 400 μg H37Ra Mycobacterium tuberculosis (DIFCO). Bordetella pertussis toxin (300 ng; Sigma Aldrich, Brøndby, Denmark) in 200 μl of PBS was injected intraperitoneally at day 0 and day 2. Animals were monitored daily from day 5 and scored on a 6-point scale as follows: 0, no symptoms; 1, partial loss of tail tonus; 2, complete loss of tail tonus; 3, difficulty walking; 4, paresis in both hind legs; 5, paralysis in both hind legs; and 6, front limb weakness. In adherence to ethical consideration, mice were sacrificed when they reached grade 6 or 24 h after hind legs paralysis. For each EAE experiment at least 16 mice were immunized (av. 50% of mice show EAE symptoms required to enter the experiment).

### 3. Cuprizone model

To induce demyelination, C57BL/6JBomTac female mice were fed with 0,3% cuprizone chow (TD.140805, Envigo, Indianapolis, IN, USA). Cuprizone-treated chow was provided fresh three times a week from a single food lot (Envigo) for five or six weeks. During the first week of the treatment, most animals exhibited a body weight loss less than 7% of their starting weight. On D0 of the study, the chow containing 0,3% cuprizone, was replaced with the structurally identical standard chow. Animals were treated with NeoMG or NeoMG-CM on days −2 and D0 (NeoMG), or −4, −2 and D0 (NeoMG-CM), followed by three days of feeding with standard chow to induce remyelination.

### 4. Magnetic-activated cell sorting (MACS)

Postnatal CAG::mRFP1 or C57BL/6JBomTac P3-5 neonates, or adult (12-16 weeks old) C57BL/6JBomTac female mice were anaesthetised by overdose of sodium pentobarbital and intracardially perfused with ice-cold PBS (Sigma-Aldrich). Brain and spinal cord tissues from neonates or brain tissue from adult animals were collected in PBS and a single cell suspension was generated by forcing through a 70-mm cell strainer (BD Biosciences). Cells were collected after centrifugation with 37% Percoll (Cytiva). Cells were magnetically sorted using CD11b MicroBeads (Miltenyi Biotec, 130-049-601) for positive selection of microglia. Viability of cells was distinguished by staining with trypan blue dye (Sigma-Aldrich). The purity was determined by flow cytometry as 80-90%.

### 5. Primary microglia cell culture

Cells isolated from the postnatal (P3-5) neonates or adult animals were suspended in Dulbecco’s Modified Eagle Medium: Nutrient Mixture F12 (DMEM/F12) (Gibco) supplemented with 10% FBS, 1X penicillin-streptomycin (Pen-Strep) (Invitrogen), 1X N-2 (Gibco, 17502048), 50 ng/mL interleukin 34 (IL-34) (BioLegend, 577602), and 50 ng/mL macrophage colony-stimulating factor (MCSF) (BioLegend, 576402). A seeding density of 100,000 cells/mL was maintained on 24-well culture plates (Falcon). Control wells contained only supplemented media. The cells were incubated at 37°C and 5% CO2. After 24 h of seeding, the cells were washed and the media was replaced with fresh DMEM/F12 medium containing 1X Pen-Strep and devoid of FBS, N-2, IL-34 and MCSF. Control media underwent identical manipulations. 12 h and 24 h after media replacement, the cell-free media was collected, frozen, and stored for subsequent use.

### 6. Cerebellar brain slice cultures

Cerebellar brain slices were obtained from C57BL/6J pups at P9-10, following previously established protocols [86]. The slices were cultured in MEM medium (Thermo Fisher Scientific), supplemented with 25% horse serum (Thermo Fisher Scientific), 25% Hank’s balanced salt solution (Sigma-Aldrich), 50 U/mL penicillin, 50 U/mL streptomycin, 1% Glutamax (Thermo Fisher Scientific), 12.5 mM HEPES (Thermo Fisher Scientific), and 1.45 g/L glucose (Sigma-Aldrich). Demyelination was induced by treating the slices with 0.5 mg/mL lysolecithin (Sigma-Aldrich) 4 days after culture initiation and maintaining the treatment for 16 h. Following lysolecithin-induced demyelination, the slices were allowed to recover in culture medium for 1 day. Subsequently, the slices were treated daily for 4 days with either vehicle (control primary microglia control medium, prepared as described in Methods section 5), NeoMG-CM or AdultMG-CM, each at a 1:1 ratio with brain slice culture medium. After the treatment, slices were fixed in 4% paraformaldehyde and stored in 1xPBS in 4°C for histological analyses.

### 7. Intrathecal injections

Animals were anaesthetised using 4% isoflurane gas (1000mg/g, ScanVet, Fredensborg, Denmark) and received analgesia (Temgesic 0.03mg/mL, VetViva Richter GmbH, Austria) and sodium chloride (0.09mg/mL, B. Braun Medical A/S, Frederiksberk, Denmark) s.c. To perform intrathecal injections, animals were placed on a platform, with the head angled down in 90°, enabling the direct injection of cells into the cerebrospinal fluid (CSF) through the cisterna magna. With the use of a 50 µL Hamilton syringe (Hamilton Company, Timis County, Romania) attached to a bent 30G needle, 10 µL of cell suspension, PBS (vehicle), or culture media was injected. Mice were randomized on four groups that received: (1) 10μl PBS (vehicle), (2) a suspension of total 3×10^5 microglia in 10μl PBS or (3) 10μl of media from NeoMG cell culture or (4) 10μl of control media. 0,3% cuprizone-treated C57BL/6BomTac female mice received injections with a two-day interval, before and on the day of cuprizone cessation. After the last injection, animals were further monitored by a blinded investigator and sacrificed 3 days after the last injection. Aged C57BL/6 female mice received three intrathecal injections before behavioural testing. Previously immunised C57BL/6BomTac female mice (EAE model) received one intrathecal injection upon reaching grade two (complete loss of tail tonus) of EAE. Animals were further monitored and scored for EAE symptoms daily by a blinded investigator for 4 days from cell transplantation. Recipient mice were sacrificed 0, 1, 2, 3 days after cell transplantation.

### 8. Behavioural testing

All behavioural tests were performed at the Biomedical Laboratory, University of Southern Denmark (Odense, Denmark). Barnes maze (BM) test, open field (OF) test, and novel object recognition (NOR) test were performed to assess the cognitive abilities of animals subjected to the treatment. All animals undergoing behavioural testing were maintained under standard controlled environmental conditions. Prior to the behavioural tests, animals underwent a 4-day handling period to minimise stress and anxiety levels. Behavioural tests were performed during the light phase, between 8.00-16.00. To eliminate the impact of olfactory cues, testing equipment was thoroughly cleaned with 70% ethanol after each trial involving a different animal. Until analysis, the experimenters were blinded to the treatment given to the animals in order to avoid any bias.

#### Barnes maze

The BM was conducted to evaluate spatial learning and memory [37], using rodents’ natural aversion to open spaces and stimulating escape from the maze. The test protocol included three days of learning (day 1-3), a short-term memory test (day 4), and a long-term memory test (day 11). During the learning phase, animals received four learning sessions per day for three days (12 sessions total), during which they were trained to locate a target with an escape box hidden under a specific hole. Thereafter, animals were subjected to short- and long-term memory tests (1 and 7 days after the last training session, respectively), with the escape box removed and the target hole covered similarly to others in the maze. Each trial began by placing the animal in a central, non-transparent chamber to ensure randomised starting positions. 15 seconds after turning on the aversive lights (IP44 Halogen work light, Elworks and P54 LED lamp, Profi Depot), the chamber was lifted and the mouse was free to explore the maze within a controlled time. Trials were recorded using two video cameras (Sony SSC-DC378P above platform, Sony HDR-CX230E beside platform). The recordings from the behavioural paradigms were thereafter analysed by an observer blinded to the treatment conditions and following parameters: the primary latency (time taken by the mouse to reach the target), number of primary errors and the target block (the average time spent in the maze quarter with the target hole), were recorded.

#### Open field

Open field test is commonly used to evaluate locomotor activity and anxiety-like behaviour in rodents [35]. To evaluate the animal’s behaviour, it was placed in the centre of a 45 × 45 × 45 cm non-transparent plastic box and recorded for 10 minutes. Locomotor activity and the incidents of various types of anxiety-indicating behaviour were recorded by the SMART video tracking system version 3.0 (Panlab, Barcelona, Spain) together with a video camera (SSC-DC378P, Biosite, Stockholm, Sweden).

#### Novel object recognition

The NOR test is based on the innate urge of rodents to explore new objects and allows to assess explorative activity and recognition memory. The test consists of three phases spread within three days. The OF test, performed on day 1 was a habituation phase, during which an animal explores the box, used for NOR test in following days. On the second day (NOR1, learning phase), the animal was placed in the box with two identical objects and allowed to explore them for 10 minutes. During the third day (NOR2, testing phase), performed 24 h after NOR1, one of the familiar objects was replaced with a novel one and the testing phase was performed for 10 min. The tests were recorded using a camera (SSC-DC378P, Biosite, Stockholm, Sweden) and then the amount of time spent by animals on exploring the objects was measured. Cognitive outcome was determined both by determining exploration time for each object and calculating discrimination index (DI) [36]. DI was calculated as: (Novel Object Exploration Time/Total Exploration Time) - (Familiar Object Exploration Time/Total Exploration Time) to assess the preference of the mouse to the object [87]. In this test, the more time spent exploring the novel object versus familiar one and the higher DI in NOR2 was considered as the indication of the improved recognition memory.

### 9. Tissue collection and processing

Animals were sedated with an overdose of sodium pentobarbital. Thereafter, they were intracardially perfused with ice-cold PBS. Following perfusion, brains were collected, and the hemispheres were separated. The left hemisphere was collected in TRIzol (Invitrogen, Fisher Scientific) and stored at - 80°C for RNA analysis. The right hemisphere was fixed in ice-cold 4% paraformaldehyde (PFA) (Sigma-Aldrich). Spinal cord and the right hemisphere from EAE animals sacrificed 0, 1, 2, 3 days after the intrathecal injection and fixed in ice-cold 4% paraformaldehyde (PFA) (Sigma-Aldrich). Upon fixation, tissue was cryopreserved in 30% sucrose (Sigma-Aldrich), embedded in optimal cutting temperature compound (O.C.T) (VWR Chemicals) and frozen in iso-pentane (VWR Chemicals) pre-cooled in liquid nitrogen. Frozen tissues were kept at −80°C until further analysis. Brain tissue was collected from the 0,3% cuprizone-treated animals, sacrificed 3 days after the intrathecal injection. Animals were anaesthetised as described above and decapitated. Thereafter, whole brain tissue was collected, fresh-frozen in CO_2_ and kept in −80°C until further analysis.

### 10. Immunostaining

Frozen brains from 10 weeks old C57BL/6 mice, 76-92 weeks old C57BL/6 mice injected with NeoMG or NeoMG-CM, or control animals, spinal cords from EAE-animals treated with NeoMG and PBS-injected vehicles as well as brains from CPZ-treated animals were cut on a cryostat (Leica CM3050 S, Leica Geosystems) for histology. 12-µm spinal cord sections, 16-µm brain sections were stored at −80°C on Superfrost Plus slides (Thermo Scientific) and 50-µm brain sections were collected in PBS and used for immunofluorescent staining upon sectioning.

12-µm spinal cord sections and 16-µm brain sections were used for immunohistochemistry. Sections were washed in PBS and incubated for 30 minutes with 0.2% H_2_O_2_ in methanol, and 1xPBS to block endogenous peroxidase. After several rinses with 0.2% Triton X-100 in PBS (PBST), they were incubated for 1 h in 3% bovine serum albumin (BSA) in PBS to block unspecific binding. Sections were immunostained for 1 h at room temperature (RT) with corresponding primary antibodies: goat anti-MOG (R&D systems, AF2439) or rabbit anti-Myelin PLP (Abcam, Cat. No. ab254363, 1:200) in 3% BSA in 0,2% PBST for 1h at RT before rinsing in 0,2% PBST. Subsequently, sections were incubated for 1h at RT with biotinylated goat anti-rabbit IgG (Abcam, Cat. No. ab64256, 1:2) secondary antibody in 3% BSA in 0,2% PBST. Following secondary antibody incubation, the sections were rinsed with 0,2% PBST and incubated for 1h at RT in streptavidin-horseradish peroxidase (HRP) conjugate (Cytiva, Cat. No. RPN1231V, 1:200) in PBST with 3% BSA. Thereafter, sections were rinsed in 0,2% PBST and PBS, and staining was developed with 3,3′-Diaminobenzidine (DAB) (Sigma-Aldrich) in 0,03% H_2_O_2_ in 1X PBS for 10 minutes. Sections were then rinsed in PBS, dehydrated in increasing ethanol concentrations, mounted, and visualized on an Olympus CX41 microscope. Stained sections were captured at 4× magnification.

Immunofluorescent stainings were performed on fifty-µm brain sections and twelve-µm spinal cord sections. TSA kit (Invitrogen, Cat. No. B40923) was used for amplification of the RFP signal in the brain sections. Brain sections were incubated 30 min in 10% methanol (Supelco) with H_2_O_2_ (TSA kit, Invitrogen) in PBS to block endogenous peroxidase, followed by rinsing in 0.2% PBST and incubation in 1% PBST for 30 min at RT. Thereafter, to block unspecific binding, sections were incubated in a blocking buffer (TSA kit, Invitrogen) for 90 min at RT. Next, sections were incubated with corresponding primary antibodies: rabbit-polyclonal to green fluorescent protein (GFP, with specificity against RFP) (1:200, Abcam, Cat. No. ab6556) and rat-anti-mouse MAC1 (1:200, Bio-Rad, Cat. No. MCA711) in 3% BSA in 0,2% PBST for 2h at RT. Following incubation, sections were rinsed with PBS and incubated with secondary antibodies: donkey-anti-rat Alexa Fluor 488 (1:200, Invitrogen, Cat. No. A21208) and poly-HRP-conjugated goat anti-rabbit secondary antibody (1:200, TSA kit, Invitrogen) for 1h at RT and overnight at 4°C. The following day, sections were rinsed in PBS and incubated 4.5 minutes in the tyramide solution (TSA kit, Invitrogen) according to the manufacturer’s protocol. After stopping the reaction, sections were rinsed in PBS and mounted. Sections were visualised with a Olympus FV1000 MPE confocal microscope 20x and analyzed with FV10-ASW 4.06 software.

12-µm spinal cord sections from EAE animals were washed in PBS and incubated for 10 min in ice-cold acetone. After repeated rinses with 1% Triton X-100 in PBS (PBST), they were incubated for 1h in 3% BSA in 1% PBST to block nonspecific binding. Next, sections were incubated overnight at 4°C with corresponding primary antibodies: rabbit anti-IBA1 (Cat. No. 019-19741 Wako) and rat anti-CD45 (Cat. No. 550539; BD). Following primary antibody incubation, the sections were washed with PBST and incubated for 1 h with the appropriate secondary antibody: donkey anti-rabbit 488 (Cat. No A21206; Invitrogen), donkey anti-rat Alexa 594 (Cat. No A21209; Invitrogen), and goat anti-rabbit Alexa 647 (Cat. No A21245; Invitrogen). Following secondary antibody incubation, sections were washed with PBS and incubated for 10 min at RT in DAPI (Invitrogen). Next, sections were rinsed in PBS and mounted using DAKO fluorescence mounting medium (Agilent Technologies). Stained sections were then visualised with an Olympus CX41 microscope.

Immunofluorescent stainings for myelin basic protein (MBP) and neurofilament (NF) were performed on cerebellar brain slices on membrane inserts. After a PBS wash, slices were blocked for 1 h at room temperature in 5% normal horse serum and 0.3% Triton X-100 in PBS. Primary antibodies-rat anti-MBP (1:250, Bio-Rad, Cat. No. MCA409S) and rabbit anti-NF (1:1000, Abcam, Cat. No. ab8135) were diluted in blocking buffer and applied overnight at 4°C. Following PBS washes, slices were incubated for 2 h at RT with Alexa Fluor-conjugated secondary antibodies: goat anti-rabbit 488 (1:500, Invitrogen, Cat. No. A11008) and goat anti-rat 555 (1:500, Invitrogen, Cat. No. A21434). After final washes, slices were mounted, covered with mounting medium and a coverslip, and subsequently imaged using a ZEISS LSM 900 Confocal Microscope.

The Luxol fast blue (LFB) staining was performed on 16-µm coronal brain sections. Sections were incubated with LFB solution ((Luxol fast blue (S3382, Sigma), 95% Ethanol, water, acetic acid) at 60°C for 2 h. After rinsing in 95% Ethanol and distilled water, sections were treated with 0.05% lithium carbonate solution for 5 sec and 70% ethanol, and then washed in distilled water. The process was repeated until the contrast between grey and white matter was clearly distinguished. Next, sections were dehydrated in increasing ethanol concentrations, mounted and captured at 4× magnification using an Olympus CX41 microscope. Mean LFB staining intensity was measured using NIH ImageJ software. Brains from 4-6 mice per group and 12 sections from each brain were analysed.

### 11. Stereology

Image processing and analyses were done using NIH ImageJ software (v.2.16.0/1.54p).

Demyelination in SC of EAE mice was measured as mean fraction area with a loss of MBP staining. SC tissue from 3-4 mice per group and 12 sections from each mouse were analysed. The outline of the spinal cord section was drawn manually using the Freehand selections tool from the Toolbar menu. Measurements of the demyelinated area were taken based on the area selection in the image (Analyze>Measure). Mean fraction area for demyelination along the spinal cord was calculated as: sum of demyelinated area*100/total area. Brains from 2-4 mice per group and 4 sections from each mouse were analysed.

Demyelination in coronal sections of CPZ-treated animals was measured as loss of LFB or PLP staining in CC using a semiquantitative method of the mean intensity values. The LFB or PLP intensity signal profile in CC was normalised by the background intensity in (Anterior cingulate area, ventral part, layer 1 (ACAv1)). Brains from 4-8 mice per group and 10-12 sections from each mouse were analysed between bregma −0,180 and −0,955. Evaluation of myelination was performed by comparing the level of staining intensity signal in analysed sections to the staining intensity signal in unmanipulated 12 weeks old C57BL/6 brain sections (considered as fully myelinated).

### 12. Electrochemiluminescence analysis for protein expression

To compare TNF, TNFR1, and TNFR2 concentrations between the NeoMG and AdultMG, media from NeoMG and AdultMG cultured for either 12 or 24 hours was analyzed with electrochemiluminescence assays as previously described (Raffaele et. al, 2024). The protein levels of TNF, TNFR1, and TNFR2 were determined in the media samples, using the V-Plex Mouse TNF-alpha assay, the U-Plex Mouse TNFRI assay, and the R-PLEX Mouse/Rat TNFRII Assay (all from Mesoscale Discovery, #K152QWD, #1520VK, #K150ZSR). A SECTOR Imager 6000 Plate Reader (Mesoscale Discovery) was used to analyze all the plates. The lower levels of detection were calculated as 2.5 standard deviations above the blank calibrator, and were for this study as follows: TNF = 0.00011 pg/ml, TNFR1 = 0.533 pg/ml; TNFR2 = 0.189 pg/ml.

### 13. Mass Spectrometry for proteomics analysis

Proteomic analysis of CM was performed using liquid chromatography-tandem mass spectrometry (LC-MS/MS). CM samples were lysed in 5% sodium dodecyl sulphate (SDS), 10 mM tris(2-carboxyethyl) phosphine (TCEP), 50 mM Triethylammonium bicarbonate (TEAB) and shaken at RT for 5 minutes at 1000 rpm, followed by boiling at 95°C for 5 min at 500 rpm and DNA digestion with Benzonase at 37°C for 15 minutes. Then samples were alkylated with 20mM iodoacetamide for 1 h at 22°C. Protein concentration was determined using EZQ protein quantitation kit (Invitrogen) as per manufacturer instructions. Protein isolation and clean up was performed using S-TRAP™ (Protifi) columns followed by digestion with trypsin at 1:20 ratio (enzyme:protein) for 2 h at 47°C. Digested peptides were eluted using 50 mM ammonium bicarbonate, followed by 0.2% aqueous formic acid and 50% aqueous acetonitrile containing 0.2% formic acid. Eluted peptides were dried down overnight followed by resuspension in 1% formic acid and injection on the Orbitrap Exploris Mass Spectrometer (Thermo Fisher).

### 14. Mass spectrometry for lipidomic analysis

Lipidomic analysis of cells or their CM was performed using liquid chromatography‒electrospray ionization tandem mass spectrometry (LC‒ESI‒MS/MS). Cell pellets or cell culture medium were reconstituted in 700 μl of 1x PBS with 800 μl of 1 N hydrochloric acid:methanol (MeOH) 1:8 (v/v), 900 μl of chloroform, 200 μg/ml of the antioxidant 2,6-di-tert-butyl-4-methylphenol (BHT; Sigma‒ Aldrich), and 3 μl of SPLASH LIPIDOMIX Mass Spec Standard (#330707; Avanti Polar Lipids, United States, Alabaster). Next, samples were centrifuged and vortexed, followed by collection of the lower organic fraction. The fraction was then evaporated with a Savant Speedvac spd111v (Thermo Fisher Scientific). The remaining lipid pellet was reconstituted in 100% ethanol. Lipid species were analyzed by LC‒ESI‒MS/MS on a Nexera X2 UHPLC system (Shimadzu, Japan, Kioto) coupled with a hybrid triple quadrupole/linear ion trap mass spectrometer (6500+ QTRAP system; AB SCIEX, The Netherlands, Nieuwerkerk aan den IJssel). Chromatographic separation was performed on an XBridge amide column (150 mm × 4.6 mm, 3.5 μm; Waters) maintained at 35 °C using mobile phase A [1 mM ammonium acetate in H_2_O-acetonitrile 5:95 (v/v)] and mobile phase B [1 mM ammonium acetate in H_2_O-acetonitrile 50:50 (v/v)] in the following gradient: (0–6 min: 0% B → 6% B; 6–10 min: 6% B → 25% B; 10–11 min: 25% B → 98% B; 11–13 min: 98% B → 100% B; 13–19 min: 100% B; 19–24 min: 0% B) at a flow rate of 0.7 ml/min, which was increased to 1.5 ml/min from 13 min onward. Sphingomyelin (SM), cholesteryl esters (CE), ceramides (CER), dihydroceramides (DCER), hexosylceramides (HCER), were measured in positive ion mode with precursor scans of 184.1, 369.4, 264.4, 266.4, 264.4, and 264.4, respectively. Tri-, di-, and monoacylglycerides (TGs, DGs, and MGs) were measured in positive ion mode with a neutral loss scan for one of the fatty acyl moieties. Phosphatidylcholine (PC), lysophosphatidylcholine (LPC), phosphatidylethanolamine (PE), lysophosphatidylethanolamine (LPE), phosphatidylglycerol (PG), phosphatidylinositol (PI), and phosphatidylserine (PS) were measured in negative ion mode by fatty acyl fragment ions. Lipid quantification was performed by scheduled multiple reaction monitoring (MRM), and the transitions were based on neutral losses or typical product ions as described above. The instrument parameters were as follows: curtain gas = 35 psi; collision gas = 8 a.u. (medium); IonSpray voltage = 5500 V and −4500 V; temperature = 550 °C; ion source gas 1 = 50 psi; ion source gas 2 = 60 psi; declustering potential = 60 V and −80 V; entrance potential = 10 V and −10 V; collision cell exit potential = 15 V and −15 V. Peak integration was performed with MultiQuantTM software version 3.0.3. Lipid species signals were corrected for isotopic contributions (calculated with Python Molmass 2019.1.1) and were quantified based on internal standard signals and adhered to the guidelines of the Lipidomics Standards Initiative. Only the detectable lipid classes are reported in this manuscript.

### 15. RNA isolation

Total RNA was extracted from brain tissue using TRIzol (Invitrogen, Waltham, MA, USA) according to the manufacturer’s protocol. RNA quality was verified with Fragment Analyzer, with all samples showing RQN values ≥ 7. Libraries were prepared using NEBNext rRNA Depletion Kit v2 and NEBNext Ultra Directional RNA Library Prep Kit (Illumina) and sequenced on a Illumina platform (NovaSeq 6000). Paired-end reads were aligned to the mouse reference genome (GRCm39, June 2020) with HISAT2, yielding 20–40 million mapped reads per sample. Differential expression analysis was performed in R using DESeq2 within the Bioconductor framework.

### 16. Proteomic and transcriptomics analysis

Proteins detected in all replicates (4 out of 4 for NeoMG or 3 out of 3 for AdMG) in at least one experimental group were retained. Contaminant proteins present in empty media controls were removed unless exhibiting at least a 10-fold increase in abundance in experimental samples. Proteins completely missing in one experimental group (MNAR) were imputed using the MinProb method, while partially missing proteins (MAR) underwent k-nearest neighbors (kNN) imputation (k = 10). Limma (10.1093/nar/gkv007) was used to identify differentially abundant proteins between groups (NeoMG vs. AdMG). For transcriptomics, DESeq2 (10.1186/s13059-014-0550-8) was used for studying the differential transcript abundance. Adjusted p-values were computed using the Benjamini– Hochberg/FDR method, with significance thresholds set at adjusted p ≤ 0.05. GO enrichment analysis (biological processes) was conducted using clusterProfiler (org.Mm.eg.db annotation, adjusted p ≤ 0.05). Redundant GO terms were reduced using the RRVGO package, employing semantic similarity matrices (“Rel” method) to identify representative parent terms. The interactome was computed using Biogrid (10.1002/pro.3978). Additional enrichment was produced by using the Monarch initiative (10.1093/nar/gkw1128), KEGGS (10.1093/nar/gkv1070), Wikipathways (10.1093/nar/gkad960) and Interpro (10.1093/nar/gkae1082) datasets. To contextualize the putative rejuvenation, our dataset was compared to the dataset of wildtype C57BL/6 samples from MODEL-AD (GSE168137). All statistical analyses and visualizations were performed using R (R Core Team (2025). R: A Language and Environment for Statistical Computing. R Foundation for Statistical Computing, Vienna, Austria. <https://www.R-project.org/>).

### 17. Lipidomic analysis

Lipids detected in at least 70% of samples in at least one group were retained. Contaminant lipids present in empty media controls were subtracted. Missing data were imputed using k-nearest neighbors (kNN) imputation. Batch effect was corrected by ComBat function from the sva package. Data were log2-transformed and Limma was used to identify differentially abundant lipids between groups (NeoMG vs. AdMG). Adjusted p-values were computed using the Benjamini–Hochberg method, with significance thresholds set at adjusted p ≤ 0.05 and |log2 Fold Change| ≥ 1. All statistical analyses and visualizations, volcano plots, were performed using R (v4.3.x) or GraphPad Prism (v10.1.0).

### 18. Statistical analysis

All statistical analyses were performed using GraphPad Prism (v10.1.0) and R (v4.4), as appropriate. One-way or two-way ANOVA with Tukey’s or Fisher’s LSD post hoc test, or Kruskal–Wallis with Dunn’s correction, was used for group comparisons. For analyses combining the 5- and 6-week CPZ groups, data were evaluated using a general linear model with Treatment and Duration as fixed factors. In the absence of a significant Treatment × Duration interaction, durations were pooled for presentation, and p-values reported from the model were adjusted for Duration. Lipidomic and proteomic data were analyzed using the limma package with Benjamini–Hochberg correction (FDR < 0.01). Data are shown as mean ± SEM. Significance thresholds: *P* < 0.05 (\**), < 0.01 (**), < 0.001 (\*\*\**), < 0.0001 (****).

## Author Contributions

AW conceptualised the study and provided financial support, AW, GL designed the research methodology. GL, ESS, ABB, DRu, MA, KNN, TC, LVB, SGSV, FM, and AW performed the experiments. GL, AW, ESS, ABB, DRu, MA, KNN and MM analysed the data. DRa performed the multi-omics analysis. AL designed and performed the proteomic experiment and contributed to proteomic data analysis. JH supervised the lipidomic and organotypic slice culture experiments and provided input on lipidomic data analysis and interpretation. KLL provided access to behavioural facilities and advised on behavioural tests and TNF data interpretation. GL and AW with support of DRa wrote the manuscript. All authors read and approved the manuscript.

## Acknowledgments

We acknowledge Dina Stær Arengoth for her valuable contributions to animal care. Technical assistance with the cerebellar slice culture experiments was provided by Leen Timmermans, and mesoscale assays were performed by Susanne Petersen. We thank Bente Finsen for providing access to equipment used in the BM test. Image acquisition was carried out at the Danish Molecular Biomedical Imaging Center (DaMBIC), University of Southern Denmark, supported by the Novo Nordisk Foundation (NNF) (grant agreement number NNF18SA0032928). Schematics of the experimental setup were created using BioRender.

