## Supplementary Fig. S1 for "Neonatal Microglia and Their Secretome as Mediators of Brain Repair"

### Supplementary Data Figures

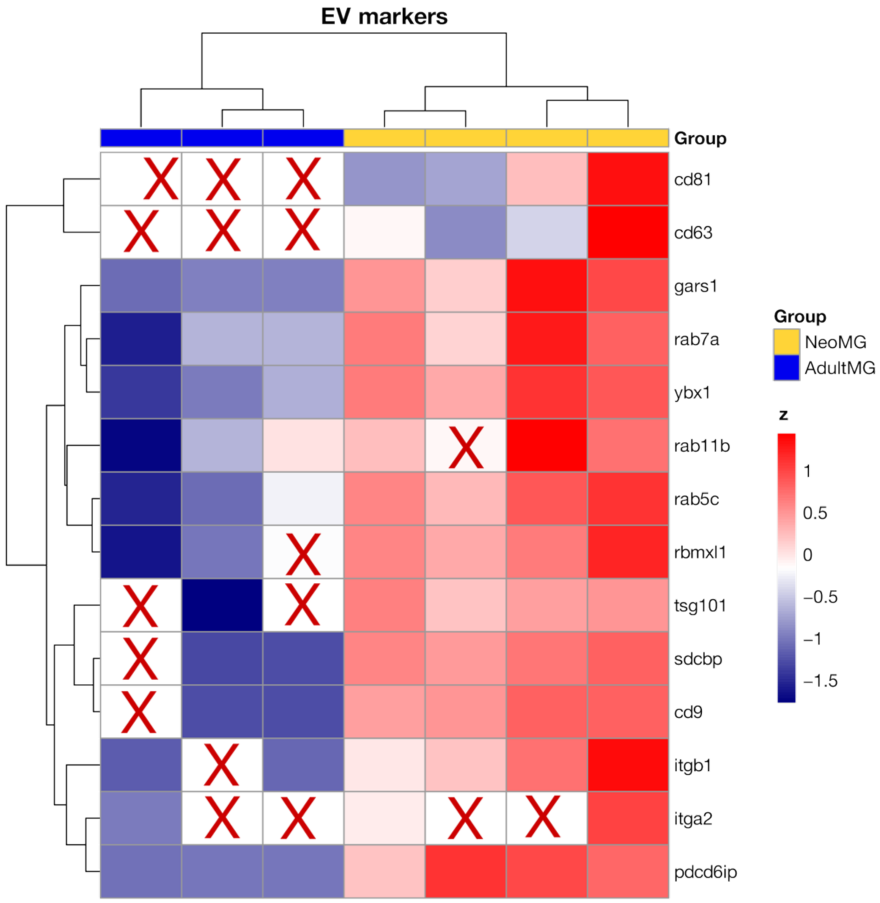


*Fig. S1. Heatmap depicting a set of EVs-related proteins [85] in NeoMG and AdultMG. Protein expression levels per sample are shown relative to the average expression across all samples. The colour key corresponds to row Z-scores, where red indicates higher expression and blue indicates lower expression.*
